# Structure and dynamics of a multidomain ligand-gated ion channel revealed under acidic conditions

**DOI:** 10.1101/2025.02.27.640520

**Authors:** Olivia Andén, Urška Rovšnik, Marie Lycksell, Marc Delarue, Rebecca J Howard, Erik Lindahl

## Abstract

Pentameric ligand-gated ion channels are critical mediators of electrochemical signal transduction across evolution, including key targets of biophysical studies and therapeutic development. However, the intrinsically allosteric, polymodal, modular nature of gating in this protein family presents persistent challenges to biophysical characterization. The bacterial channel DeCLIC constitutes a provocative model system for structure, function, and dynamics in this family, including a modulatory N-terminal domain (NTD). Previous closed structures of DeCLIC support a rationale for its inhibition by calcium via a site of conserved relevance in other family members; however, mechanisms of gating in DeCLIC and properties of its open state have remained unclear. Here we integrated structure-function methods including cryogenic electron microscopy (cryo-EM), molecular dynamics simulations and small-angle neutron scattering under acidic conditions to characterize a previously unreported conformation of DeCLIC with a fully hydrated pore. In contrast to previous structures, the low-pH open conformation captured by cryo-EM was stable and permeable in simulations, and consistent with solution-phase behavior as measured by small-angle scattering. We further captured an alternative closed state of the channel, evidently promoted by depletion of modulatory calcium at low pH, exhibiting dynamic rearrangements in the N-terninal domain. The expanded-pore structure evidently corresponds to a functional open state of DeCLIC, while calcium-site and NTD dynamics drive channel closure, providing a detailed template for biophysical characterization of modulatory mechanisms in ligand-gated ion channels and related systems.

**Significance Statement:** The bacterial protein DeCLIC is a provocative representative of the pentameric ligand-gated ion-channel family. Like its eukaryotic relatives, DeCLIC is sensitive to external calcium, and incorporates a possibly disordered modulatory domain. But as for many ion channels, the details of DeCLIC activation have remained unclear, as no stable open state has been characterized. Combining biophysical methods, including cryogenic electron microscopy, molecular dynamics simulations, and small-angle neutron scattering, allows us to capture, validate, and characterize an evidently open state of DeCLIC, as well as a closed state with a disordered modulatory region. Details of these structures, their function, and their dynamics allow us to propose a mechanism for DeCLIC gating, as well as a generalized scheme for the larger protein family.

## Introduction

Pentameric ligand-gated ion channels (pLGICs) constitute a family of membrane proteins expressed in numerous eukaryotic and prokaryotic organisms [1]. These receptors open an intrinsic ion-conducting pore in response to chemical stimuli, with possible ligands ranging from neurotransmitters to protons [2]. In mammals, pLGICs are important contributors to the nervous system, as they facilitate the conversion of chemical to electrical signals at synapses [2]. Dysfunction in pLGICs is connected to several neurological disorders including Alzheimer’s, epilepsy, hyperekplexia, and addiction [3]. Accordingly, these receptors are targets for therapeutic agents including benzodiazepines, steroids, and general anesthetics [4]. A variety of pLGICs are also subject to modulation by environmental factors including extracellular calcium (Ca^2+^) [5–7] and pH [8–10].

All known pLGICs adopt a similar architecture, with an extracellular or periplasmic domain (ECD) consisting of 10 β-strands (β1-β10) [11], including the characteristic Cys-loop (in eukaryotes) or Pro-loop (in prokaryotes) between β6-β7 [12], and a transmembrane domain (TMD) consisting of four α-helices (M1-M4) [11]. Eukaryotic pLGICs also have an intracellular domain (ICD) of varying length between the M3 and M4 helices [13], while prokaryotic pLGICs can contain an additional N-terminal domain (NTD) [1]. These peripheral domains are thought to modulate channel activity, along with a host of environmental stimuli including allosteric ligands, ions, and pH. Despite recent advances in structure determination, mechanisms of pLGIC permeation and modulation in many cases remain unclear. The functionally critical open state is particularly underrepresented in structural data, due in part to the rapid desensitization seen especially in eukaryotic family members [14, 15].

Recently, we have characterized fundamental structure-function properties in a prokaryotic member of the pLGIC family, derived from a *Desulfofustis* deltaproteobacterium [16]. This channel, called DeCLIC, includes the canonical ECD and TMD, plus an NTD comprising roughly half the receptor mass. Electrophysiological experiments show that DeCLIC is modulated by Ca^2+^ ions, and the channel crystallizes in either a closed or wide-open state in the presence or absence of Ca^2+^, respectively [16]. We have also used cryogenic electron microscopy (cryo-EM) and molecular dynamics (MD) simulations to propose a nonlinear ion pathway via fenestrations in the ECD that can be blocked by Ca^2+^ [17]. Furthermore, MD and small-angle neutron scattering (SANS) reveal substantial rigid-body motions of individual lobes of the NTD. Interestingly, MD and SANS data have proven inconsistent with a functional role for our initially reported wide-open X-ray structure; moreover, predominant states identified by cryo-EM are closed in both the presence and absence of Ca^2+^. These findings raise provocative questions as to the functionally relevant open state, pathway for ion conduction, and modulatory role of the NTD.

Here we describe a previously unreported open structure of DeCLIC, along with alternative closed states from the same cryo-EM grids, by preparing samples under acidic conditions (pH 5). This open state is more stable in MD simulations than our previous X-ray structure, and consistent with lipid-bilayer recordings and SANS profiles under room-temperature conditions. In combination with previous functional and structural data, we propose a mechanism for modulation by factors including pH, Ca^2+^, and NTD flexibility, offering a template for comparable mechanisms in the larger pLGIC family.

## Results

### Closed and open states in a single low-pH dataset

Previous structure-function studies of DeCLIC indicate the presence of an inhibitory binding site for Ca^2+^ at the periplasmic subunit interface, featuring at least four Glu residues [16, 17]. Given the likely sensitivity of such a coordination site to changes in pH, and the sensitivity of several related pLGICs to proton activation [27, 28], we tested whether DeCLIC might adopt an alternative conformation under acidic conditions. For DeCLIC samples prepared at pH 5 with 10 mM Ca^2+^, we observed two distinct populations, with either a constricted or expanded conformation of the pore. Processing each population individually resulted in an overall resolution of 3.1 Å for the constricted- and 2.9 Å for the expanded-pore state (Figure 1A, Supplementary Tables **??** and **??**, Supplementary Figure **??**). Local resolution in the TMD and ECD was better than in the NTD (Supplementary Figure **??**D), where some sidechain and backbone atoms could not be built with confidence, particularly for the constricted state (Supplementary Table **??**).

**Figure 1:**
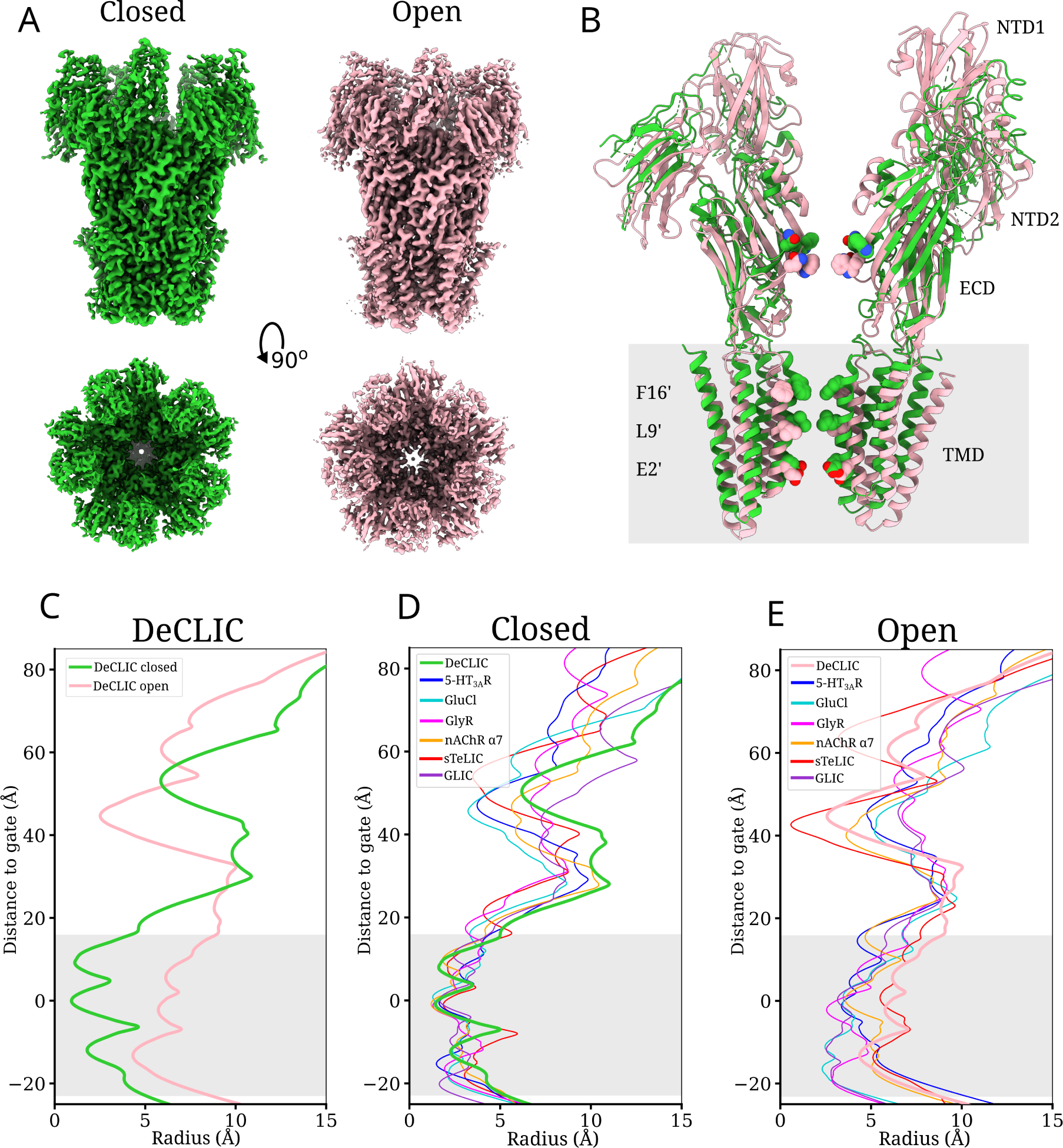
Closed and open states in a single low-pH dataset. (A) Cryo-EM reconstructions of DeCLIC in closed (lime) and open (pink) conformations, viewed from the membrane plane (*top*) and down the pore from the extracellular side (*bottom*). (B) Two subunits on opposite sides of each other are shown in ribbon, both for closed (lime) and open (pink) DeCLIC. The membrane is marked with a grey background, and individual lobes and domain contributors of the righthand subunit (NTD1, NTD2, ECD, TMD) are labeled. Constricting residues in the pore as well as the ECD are displayed as spheres and colored by heteroatom. (C) Pore profiles (including sidechains) of the closed (lime) and open (pink) structure determined at pH 5. (D) Pore profiles (including sidechains) of closed DeCLIC (lime) compared to previously reported closed structures of serotonin receptor (PDB ID: 6BE1 [18], blue), glutamate-gated chloride channel (PDB ID: 4TNV [19], cyan), glycine receptor (PDB ID: 6PM3 [20], magenta), α7 nicotinic acetylcholine receptor (PDB ID: 7KOX [21], orange), sTeLIC (PDB ID: 9EX6 [22], red), and GLIC (PDB ID: 6ZGD [23], purple). (E) Pore profiles (including sidechains) of open DeCLIC (pink) compared to previously reported open structures of serotonin receptor (PDB ID: 6DG8 [24], blue), glutamate-gated chloride channel (PDB ID: 3rif [25], cyan), glycine receptor (PDB ID: 6PM6 [20], magenta), α7 nicotinic acetylcholine receptor (PDB ID: 7KOO [21], orange), sTeLIC (PDB ID: 9F5N [22], red), and GLIC (PDB ID: 4HFI [26], purple).

We built two independent DeCLIC structures into their respective densities from this dataset. The constricted-pore structure included a TMD comparable to previous closed structures [16, 17], especially at the L554 (9’, hydrophobic gate) and F561 (16’) levels, where the radius of the pore was 1.4 Å and 1.6 Å, respectively (Figure 1B-C). This structure is hereafter referred to as closed. In contrast, the expanded-pore structure contained a 9’ radius of 5.8 Å and a narrowest constriction, at E437 (2’), of 4.4 Å (Figure 1B-C), and is hereafter referred to as open. Although the transmembrane pore profile of this open structure diverged from our previous X-ray structure of DeCLIC in a wide-open state [16], it was similar to the homologous prokaryotic channel sTeLIC [28]. Comparing our two DeCLIC structures to other homopentameric pLGICs for which both closed and open structures are available shows a notable similarity among closed structures in the transmembrane pore (Figure 1D), especially at the hydrophobic gate (9’) where the pore radius ranges only 0.7 Å (1.2 Å to 1.9 Å). Interestingly, pore profiles of the corresponding open structures vary more widely, with a radius ranging 3.2 Å (2.6 Å to 5.8 Å) at the hydrophobic gate (Figure 1E).

In the ECD, the closed DeCLIC structure contained a constriction to 6 Å formed by the sidechains of residues N405 and W407, which contracted even further to *≤*3 Å in the open structure (Figure 1B-C). Moreover, the closed structure included a distinct non-protein density in the vicinity of the principal β1-β2 loop and complementary loop F, proximal to sidechain oxygen atoms of residues E347 and E480. A superimposable density was previously observed by Ba^2+^ anomalous scattering, and recapitulated in closed X-ray and cryo-EM structures determined at pH 7 in the presence but not the absence of Ca^2+^ [16, 17, 29], and was accordingly assigned as a Ca^2+^ ion (Figure 2A-B).

**Figure 2:**
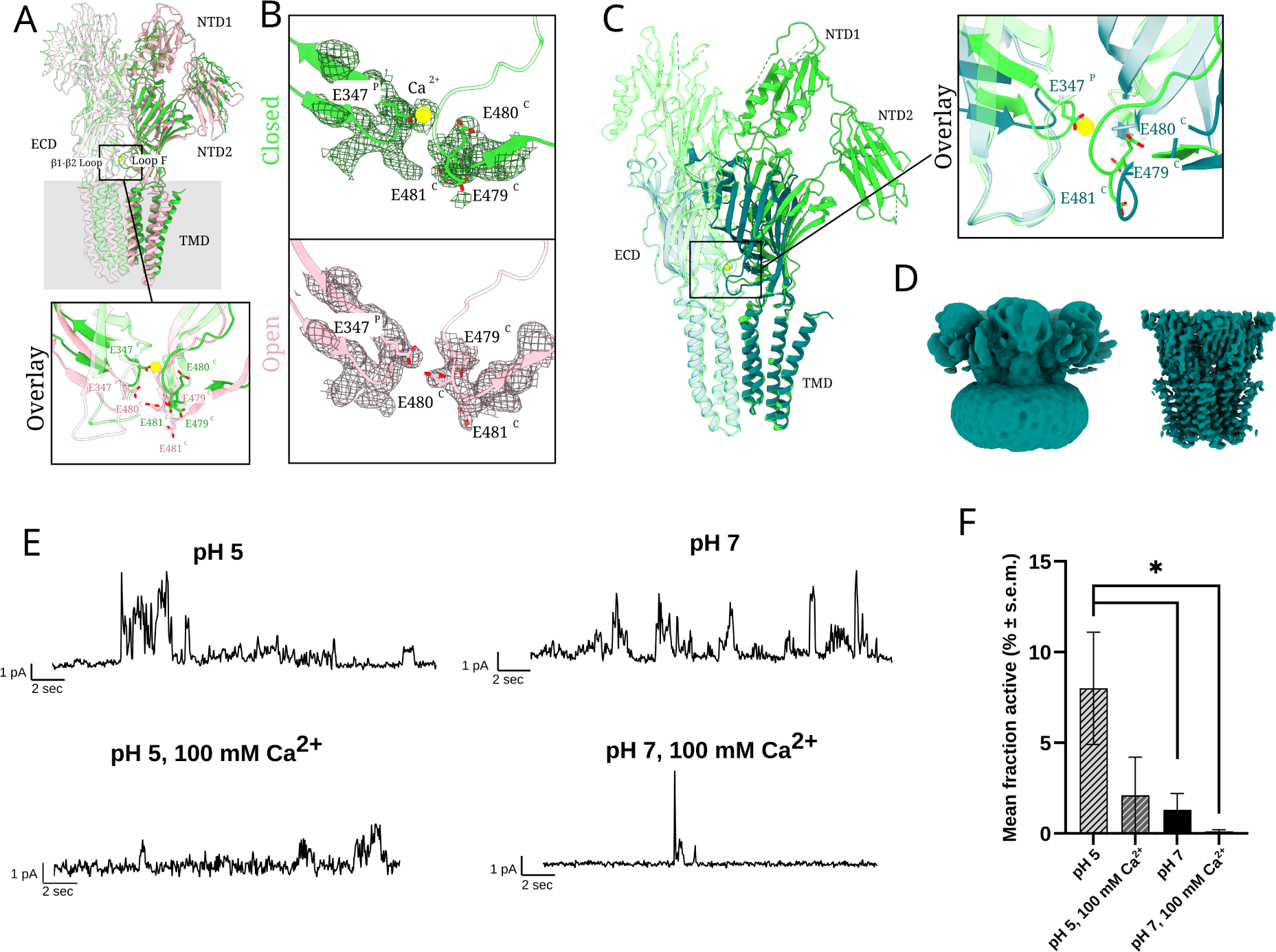
Calcium dissociation and channel opening at low pH. (A) Overlay of two adjacent subunits of closed (lime) and open (pink) DeCLIC determined at low pH with Ca^2+^. Lower inset highlights the interdomain Ca^2+^site between β1–β2 from the principal subunit (P, semitransparent) and loop F from the complementary subunit (C, opaque), showing coordinating sideshains (sticks colored by heteroatom) and Ca^2+^ (yellow). (B) Zoom views of closed (*top*, lime) and open (*bottom*, pink) DeCLIC structures as in *A*, with overlaid cryo-EM densities (mesh). (C) Overlay of two adjacent subunits of closed (lime) and partly disordered (teal) DeCLIC determined at low pH in the presence and absence of Ca^2+^, respectively. Righthand inset highlights the interdomain Ca^2+^ site as in *A*. (D) Cryo-EM reconstruction of partly disordered DeCLIC viewed at low (*left*) and high (*right*) threshold. (E) Sample traces from electrophysiology recordings in DPhyPC bilayers clamped at +50 mV at pH 5 (*left*) or pH 7 (*right*), in the absence (*top*) or presence (*bottom*) of 100 mM Ca^2+^. (F) Mean fractional recording time (± standard error of the mean) with apparent DeCLIC activity (conductance *>* 4 pS) in bilayers prepared as in *E*. In the absence of Ca^2+^, channel function increases under acidic conditions (*n* = 19–21, \**p <* 0.05 by two-tailed unpaired *t*-test).

The open structure, resolved to a comparable resolution from the same pH-5 grid, did not include this non-protein ECD density; instead, it featured substantial remodeling of Glu-containing loops in the open relative to the closed structure. On the principal side, the β1-β2 loop shifted towards the TMD (downward), displacing the sidechain of residue E347 from its position coordinating Ca^2+^ from the principal side. On the complementary side, loop F also moved away from the Ca^2+^ site, shifting towards the ECD vestibule (inward). These down-and-in motions corresponded to an overall contraction and counter-clockwise rotation of the the ECD with respect to the TMD when viewed from the periplasmic side. The open structure also exhibited rigid-body contraction of both NTD lobes compared to the closed structure, increasing the intra-subunit buried surface area between NTD1 and NTD2 by 237 Å^2^ and between NTD2 and the ECD by 196 Å^2^, possibly contributing to the improved resolution of the NTD backbone in the open state (Figure 1A-B and Supplementary Table **??**). Despite their differences in the TMD, the open-state ECD and NTD were consistent with those of our previous wide-open X-ray structure [16], supporting the generalizability of these transitions under neutral or acidic conditions.

### Calcium dissociation and channel opening at low pH

We next collected cryo-EM data at pH 5 with 10 mM ethylenediaminetetraacetic acid (EDTA) rather than Ca^2+^, to explore the latter’s role in DeCLIC modulation. We again detected a population with an open pore, resolvable to 3.5 Å, and superimposable with the open cryo-EM reconstruction described above. A second population at pH 5 in the absence of Ca^2+^ featured a TMD similar to previous closed structures, but a distinct conformation in the ECD and NTD, with the consensus refinement resolving to 4.0 Å overall (Figure 2C-D). In the ECD, although some sidechains could not be built, backbone remodeling in both the β1-β2 loop and loop F indicated an overall clockwise rotation relative to previous closed structures, opposite to the counter-clockwise rotation observed in the open state (Figure 2C). Local resolution in the NTD was particularly poor, likely attributable to a range of distinct conformations that averaged out in a majority of 3D classes; this structure is hereafter referred to as partly disordered.

To further investigate the apparent heterogeneity of the NTD in this population, we trained a cryo-DRGN [30] latent variable model using the partly disordered particles. By *k*-means clustering of the resulting latent space, we were able to refine multiple densities with distinct conformations of the NTD, and to rigid-body fit individual NTD lobes in at least two conformations (Supplementary Figure **??**). Interestingly, the resulting models were reminiscent of previous MD snapshots and small-angle scattering behavior [17], including displacement of individual NTD lobes outward and/or downward from their positions in higher-resolution structures.

Given that both our cryo-EM datasets under acidic conditions included an open state of DeCLIC, we also sought to characterize the function of this channel at low pH. Although we previously documented DeCLIC currents in *Xenopus* oocytes upon Ca^2+^ depletion at neutral pH [16], equivalent recordings at low pH were compromised by high, unstable background and poor cell viability. We obtained more consistent results from purified DeCLIC in planar lipid bilayers, where in the absence of Ca^2+^, currents were apparent at pH 5 with greater fractional activity than at pH 7 (Figure 2E-F, Supplementary Figure **??**, Supplementary Table **??**). Transient currents were also observed in the presence of Ca^2+^, though activity was near completely suppressed in 100 mM Ca^2+^ at pH 7. No currents were observed under equivalent conditions in bilayers without DeCLIC (Supplementary Figure **??**E). Thus, our functional as well as structural data demonstrate enhanced DeCLIC opening at pH 5.

### MD simulations support a stable, conductive open state

We next conducted all-atom MD simulations to test the stability of our DeCLIC structures. Below neutral pH, the fractional protonation of acidic amino-acid residues is expected to increase; therefore, we calculated the local pKa of all 76 Asp and Glu residues in each DeCLIC chain in both closed and open structures [31, 32], and fixed their protonation states to approximate pH 5 (Supplementary Figures **??**, **??** and **??**). In solution, the sidechain pKas of Asp and Glu are below 5, such that both amino acids are primarily deprotonated even at pH 5. However, for 30/76 such residues, conditions in the DeCLIC interior raised the local pKa, such that they were simulated in neutral form in at least one condition. Notably, 12 Asp and Glu residues were predicted to protonate at pH 5 in a state-dependent manner consistent with acidic pH-gating: local pKa at these positions increased by at least 0.5 units in the open versus closed states, supporting a link between side-chain protonation and channel activation (Supplementary Figure **??**). These prospective proton sensors were distributed across the NTD and ECD, suggesting involvement of multiple sites in channel activation; interestingly, they included E347, E479, E480, and E481, the four Glu residues in the interfacial Ca^2+^ site (Figure 2A). 18 other residues were predicted to protonate with limited or inverted preference for the open state, unlikely to drive proton gating (Supplementary Figure **??**).

Both the closed and open structures retained their respective pore conformations throughout quadruplicate 1-µs simulations (Figure 3A and B, Supplementary Figure **??**). For the partly disordered state obtained in the absence of Ca^2+^ (Figure 2C and D), lack of a complete model in the periplasmic regions precluded comparable simulations. Although we initially included the resolved Ca^2+^ ions in simulations of the closed structure, they frequently dissociated from the protein (Supplementary Figure **??**). Such dissociation was not observed in previous simulations at neutral pH [17], consistent with increased affinity upon deprotonation of Ca^2+^-binding residues. Dynamic changes were also observed in the NTD, where individual NTD1 and NTD2 lobes underwent rigid-body motions relative to the canonical ECD-TMD region, largely producing snapshots with extended NTDs; interestingly, these motions were lower in acidified (protonated) versus neutral (deprotonated) simulations of the same state (Supplementary Figures **??** and **??**). Along with the generally improved resolution of our structures at pH 5 versus previous structures at pH 7 [17], these results support a stabilizing effect of acidification particularly on the NTD, at least in the ordered closed and open states (Figure 2A and B).

**Figure 3:**
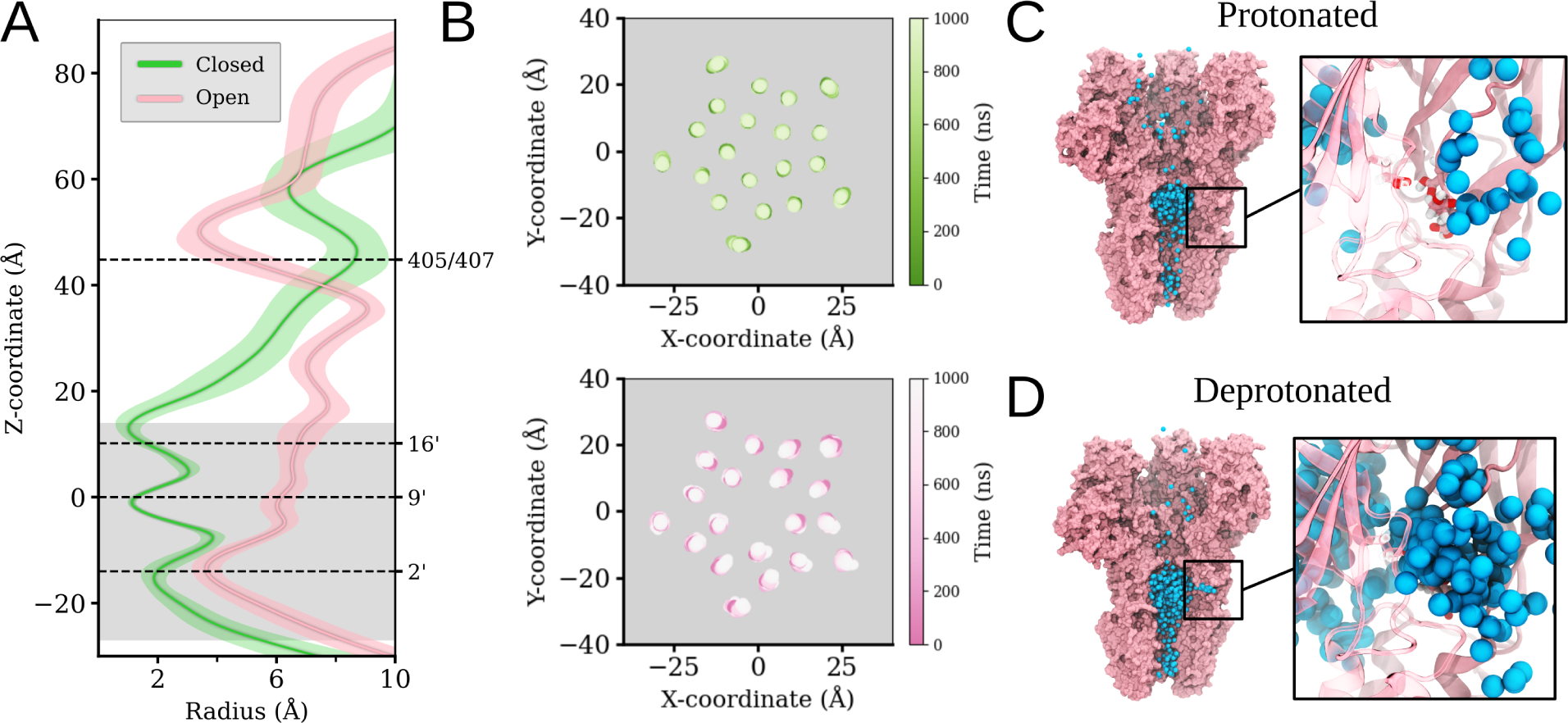
MD simulations support a stable, conductive open state. (A) Pore profiles (mean ± standard deviation) from four 1-µs replicate simulations each of closed (lime) and open (pink) DeCLIC structures determined under acidic conditions, with protonation states set to approximate pH 5. (B) Center-of-mass positions of DeCLIC transmembrane helices, viewed from the extracellular side, over simulations of the closed (*top*, lime) and open (*bottom*, pink) structures. (C) Render from a single simulation of the open structure set to approximate pH 5 (protonated), showing one representation per ns of three sodium ions (blue) which diffuse through the pore. Inset shows a zoom view of the Ca^2+^ site, with all sodium ions in the vicinity of the site across the same trajectory. The proximal subunit is hidden for clarity. (D) Render and inset as in *C* from a single simulation of the open structure set to approximate pH 7 (deprotonated).

In both neutral (deprotonated) and acidic (protonated) simulations of the open cryo-EM structure, ions could be observed spontaneously permeating the pore (Supplementary Figures **??** and **??**). In addition to traversing the length of the protein’s central axis, some sodium ions transited between the transmembrane pore and aqueous medium via the intersubunit site between the β1-β2 loop and loop F, particularly under deprotonated conditions (Figure 3C and D). Sodium ions permeated more frequently than chloride in deprotonated systems (Supplementary Figure **??**), consistent with the presence of acidic residues both in the interdomain site and in the presumed selectivity filter at the intracellular end of the transmembrane pore [16]. In protonated simulations, some chloride permeation was also observed (Supplementary Figure **??**); however, anion permeability is likely overestimated relative to functional conditions, given that our simulations fixed protonation states corresponding to acidic conditions in the intracellular as well as periplasmic compartments.

### Solution-phase occupancy of the open state determined by small-angle scattering

To test the validity of our open cryo-EM structure under room-temperature, solution-phase conditions, we next collected small-angle scattering data from DeCLIC at pH 5 in the presence and absence of Ca^2+^ (Supplementary Tables **??** and **??**, Supplementary Figure **??**). We used neutron scattering and deuterated detergent to avoid contributions from the micelle [33], and employed a paused-flow procedure including an inline size-exclusion chromatography column (SEC-SANS) previously shown to detect conformational changes in pLGICs [34]. As calculated by Guinier analysis, the radius of gyration (*R*g) was slightly larger in the absence versus presence of Ca^2+^ (51.6 ± 0.2 Å versus 50.7 ± 0.2 Å) (Figure **??**, Supplementary Table **??**), consistent with at least a partial population with an extended NTD. Comparing SANS data to curves predicted from our pH-5 cryo-EM structures (Figure 4A, Supplementary Table **??**) indicated a better fit to the open than closed structures in both the presence (χ^2^ 6.8 to open, 47 to closed) and absence of Ca^2+^ (χ^2^ 16 to open, 28 to closed). Pairwise distance distributions from both pH-5 SANS profiles were also comparable to predictions from the open structure, while the closed structure better approximated SANS curves collected previously at pH 7 with Ca^2+^ (Figure 4B) [17].

**Figure 4:**
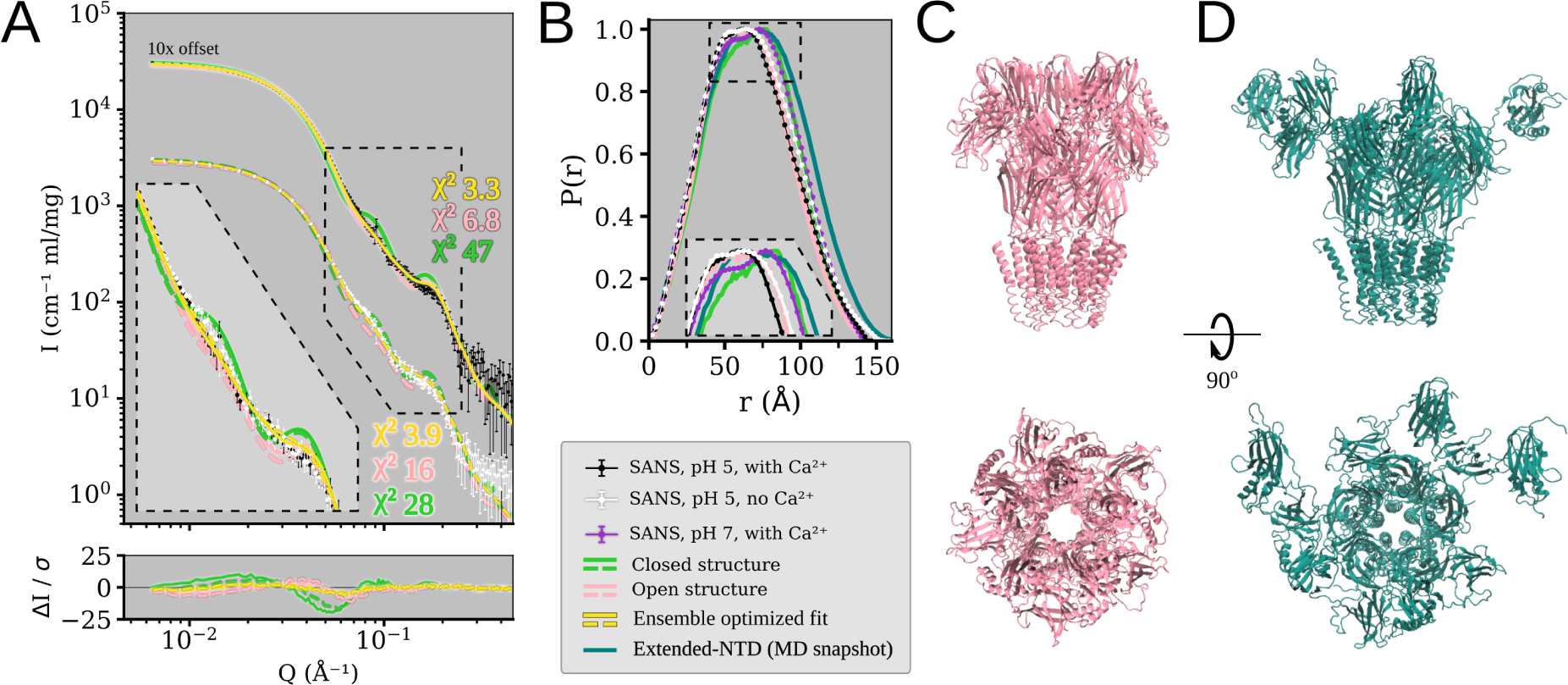
Solution-phase occupancy of the open state determined by small-angle scattering. (A) Small-angle neutron scattering profiles from DeCLIC at pH 5 with Ca^2+^ (black, x10 off-set intensity) and without Ca^2+^ (white), each plotted with theoretical scattering profiles for the pH-5 closed (lime) and open (pink) structures and an ensemble optimization fit (yellow). χ^2^ values indicate goodness of fit for each theoretical profile to the SANS data. Inset shows the main feature of the profile, without the intensity off-set. Lower panel shows the error-weighted residual of the comparisons between experimental and theoretical scattering profiles. (B) Pair-distance distributions from small-angle scattering data at pH 5 with Ca^2+^ (black), pH 5 without Ca^2+^ (white), and pH 7 with Ca^2+^ (purple) [17], from predictions of closed (lime) and open (pink) structures, and from the ensemble optimization model (teal). (C) Simulation snapshot selected by ensemble optimization in an apparent open state. (D) Simulation snapshot selected by ensemble optimization in an extended-NTD conformation.

Given that both low-pH cryo-EM datasets yielded both open and closed structures, we hypothesized that multiple states coexist in solution. Fitting linear combinations of closed and open structures marginally improved correspondence to the SANS profiles (Supplementary Figure **??**A and B), with the best fits incorporating 92% (χ^2^ 6.5) or 71% (χ^2^ 13.9) open-structure contributions under Ca^2+^ and Ca^2+^-free conditions, respectively. Including an alternative-NTD conformation from cryoDRGN analysis of the partly disordered state (Supplementary Figure **??**) further improved the fits. Indeed, the best of these multi-component fits contained only contributions from open and alternative-NTD models (91% open and 9% alternative-NTD for the Ca^2+^ dataset, χ^2^ 5.6; 79% open and 21% alternative-NTD for the dataset without Ca^2+^, χ^2^ 11.6) (Supplementary Figure **??**C-F).

### Dynamic ensemble of functional states implicated by integrating MD and SANS

To better account for dynamic rigid-body motions of NTD lobes observed in our simulations, which might be poorly represented in static structures, we calculated theoretical SANS curves for frames sampled every 10 ns in our MD trajectories. Comparing these to our experimental scattering data (Supplementary Figure **??**) identified snapshots with improved goodness of fit to each dataset, relative to static models. For the Ca^2+^ dataset, the best-fitting MD snapshot came from pH-5 simulations of the open structure, with an asymmetric NTD (χ^2^ 5.4). For the Ca^2+^-free dataset, MD sampling improved goodness of fit even more, albeit with no clear preference for snapshots from closed (best χ^2^ 9.5) or open (best χ^2^ 9.8) simulations.

We further leveraged conformational sampling from MD to fit our scattering data through ensemble optimization. We first optimized ensemble fits to our pH-5 SANS data by applying the genetic algorithm Gajoe from Eom [35, 36] to a pool of simulation snapshots from this and our previous work [17]. This approach yielded good fits to all datasets (Supplementary Table **??**, Figure 4A). Ensembles optimized to our pH-5 SANS data contained snapshots from simulations of open and closed structures, in different proportions depending on the presence (82% open, 18% closed, χ^2^ 3.3) or absence of Ca^2+^ (57% open, 43% closed, χ^2^ 3.9). Notably, open snapshots in the ensembles were visibly similar to the open cryo-EM structure, with relative symmetry in the NTD; in contrast, closed snapshots consistently featured some of the most extended NTD arrangements sampled (Figure 4C-D).

To extend these insights to neutral conditions, we also applied ensemble optimization using the same MD trajectories to fit our previously published SANS curves collected at pH 7 [17]. Interestingly, the resulting ensembles contained not only open and extended-NTD models, but also snapshots similar to the consensus closed structure [16] (Supplementary Figure **??**, Supplementary Table **??**). These closed-like models were the largest contributors to ensemble fitting of the pH-7 SANS dataset with Ca^2+^, and lesser contributors to the pH 7 dataset without Ca^2+^. Extended-NTD snapshots made similar contributions in ensembles fitting either pH-7 SANS dataset. On the other hand, open-like contributions increased at pH 7 in the absence of Ca^2+^ (Supplementary Figure **??**, Supplementary Table **??**), consistent with electrophysiology recordings in oocytes [16]. Thus, MD-based fitting of SANS data produced structural ensembles consistent with several characteristics of DeCLIC structure-function, including a relatively ordered open state, especially under acidic conditions; an ordered closed state under neutral conditions, especially in the presence of Ca^2+^; and partial occupancy of a relatively disordered nonconducting state under all conditions.

## Discussion

A growing catalog of atomistic structures, enabled particularly by the cryo-EM resolution revolution, leaves critical questions as to the foundational landscape and evolutionary diversity of ligand-gated ion-channel gating. In pursuit of these gaps, our present work combined cryo-EM, MD simulations and small-angle scattering to characterize state-dependent differences in DeCLIC domain structure and dynamics. Our data are consistent with a minimal mechanism for pLGIC gating, in which the transition from closed to open corresponds to a relative rotation and contraction of the ECD and expansion of a central hydrophobic gate in the TMD (Figure 5A). DeCLIC further elaborates on this scaffold by binding Ca^2+^ in the closed-state ECD, opening at acidic pH, and including an NTD that is especially flexible in the context of a closed pore (Figure 5B). In contrast, eukaryotic pLGICs elaborate on the minimal scaffold by binding ligands in the ECD and opening upon ligand binding (Figure 5C). They also include a likely disordered ICD which thus far is largely inaccessible by cryo-EM. The minimal mechanism can be further elaborated by additional metastable states: Apart from closed and open, DeCLIC can occupy a partly disordered nonconducting state with a differentially rotated ECD and particularly flexible NTD (Figure 5B), while eukaryotic channels adopt desensitized states with an activated ECD and differentially constricted pore (Figure 5C).

**Figure 5:**
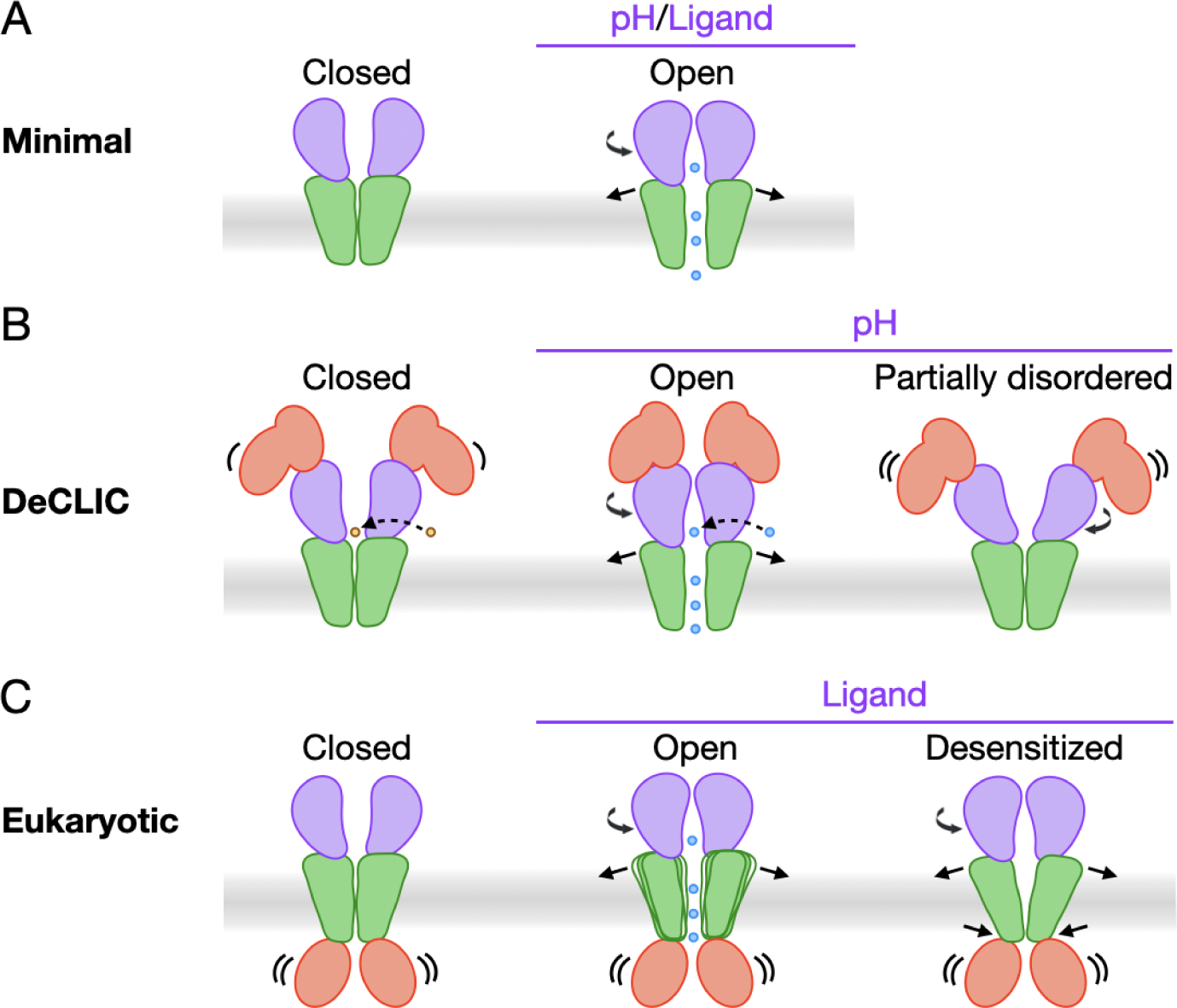
An evolving picture of pLGIC gating. (A) A minimal mechanistic scaffold for pLGIC gating, in which contraction and counter-clockwise rotation of the ECD (purple) enable opening of the TMD pore (green) and permeation of ions (blue) across the cell membrane. (B) A subtype-specific mechanism for DeCLIC gating as reported here, in which extracellular acidification, dissociation of ECD Ca^2+^ (yellow), stabilization of the NTD (red), and opening of the TMD pore (green) are associated with permeation of Na^+^ (blue) via either linear or lateral pathways. Cryo-EM, MD, and SANS experiments also show evidence of a partly disordered state featuring clockwise rotation of the ECD, dynamic displacements among NTD lobes, and a nonconducting pore. (C) A generalized gating mechanism for eukaryotic pLGICs, in which ligand binding in the ECD (purple) and opening of the TMD pore (green) enable permeation of ions (blue). A likely disordered ICD (red) may further influence gating dynamics. An additional desensitized state features a ligand-bound ECD but a nonconducting pore, distinct from the resting-closed state. Whereas closed states exhibit comparable pore profiles, open states are relatively diverse, with some eukaryotic family members substantially contracted relative to bacterial homologs like DeCLIC.

Characterizing an apparently functional open state of DeCLIC addresses an ongoing challenge in ion-channel structure-function, as open pLGIC structures remain difficult to capture. Indeed, a verified open state has not yet been reported for classical GABA(A) or nicotinic receptor subtypes expressed at neuronal synapses, where desensitization kinetics and non-physiological membrane mimetics may particularly limit open-state stability [37]. Interestingly, whereas domain arrangements and pore profiles are largely convergent among closed pLGIC structures, open structures reported thus far exhibit a range of profiles (Figure 5C). Indeed, resting-closed structures of DeCLIC have proved largely superimposable at pH 7 with or without Ca^2+^, with Ca^2+^ at either pH 7 or pH 5, in either detergent or nanodiscs [29], and are generally similar to other closed pLGICs. Conversely, the open DeCLIC pore reported here is wider than most reported in the family, similar only to those of sTeLIC and the human α7 nicotinic acetylcholine receptor [21, 28]. Despite this relative expansion, intersubunit interfaces remain tight enough to exclude membrane lipids from the channel pore, unlike our previously reported wide-open X-ray structure of DeCLIC and some recent structures of other family members [16, 38]. The open state reported here is also more stable in MD simulations than our previous wide-open X-ray structure, and consistent with SANS data under both neutral and acidic conditions. The biophysical or physiological relevance, if any, of the wide-open state remains unclear, although it recalls mixtures of apparent open and hyper-expanded states previously observed for eukaryotic Gly receptors [20]. A key experimental observation in this work was the enhancement of DeCLIC activation at low pH (Figure 5B). Whereas previous cryo-EM data collected under neutral conditions were dominated by the closed state, acidification of the sample buffer allowed us to capture a partial open population in both the presence and the absence of Ca^2+^. Recordings in Ca^2+^-free lipid bilayers were also consistent with increased function at acidic pH, and SANS profiles supported an increased proportion in the open state under acidic versus neutral conditions in the presence of Ca^2+^. Although protons were not directly observable in our cryo-EM densities, pKa calculations based on closed versus open structures predicted state-dependent neutralization of multiple acidic residues in the NTD and ECD. Among others, these included E347, E479, E480, and E481 at the ECD-TMD interface, where dramatic rearrangements were associated with Ca^2+^ dissociation and channel opening. Multidisciplinary studies of the homologous ion channel GLIC, including X-ray crystallography, cryo-EM, mutagenesis, electrophysiology, infrared spectroscopy, and constant-pH simulations [23, 39, 40], have similarly implicated multiple positions in proton gating. Interestingly, a key proton sensor in GLIC (E35) [39] aligns with DeCLIC-E347, indicating that sites of pH sensing could be partially conserved across evolution.

At acidic pH with Ca^2+^ at low pH, the open state was in apparent equilibrium with the resting-closed state, offering insights into the influence of Ca^2+^ on the pLGIC gating landscape. At acidic as well as neutral pH, the closed state of DeCLIC is capable of binding Ca^2+^ at the ECD-TMD interface (Figure 5B). Channel opening is associated with dramatic rearrangements in this region, evidently incompatible with coordinating Ca^2+^, which is absent in the open state even on a grid where the closed state is Ca^2+^-bound. Evacuation of Ca^2+^ from the domain interface also enables permeation of ions via fenestrations to and/or from the transmembrane pore. On the other hand, previous cryo-EM data under neutral conditions show a conserved backbone configuration with and without Ca^2+^ [17], indicating ECD-Ca^2+^ binding may be more a consequence than a stimulus of channel opening. Although the only non-protein densities apparent in the present Ca^2+^ structures were in the ECD, parallel work has also identified Ca^2+^ interactions in the NTD [16, 29], consistent with disordering of this domain in closed channels when Ca^2+^ is removed (see below). Thus, as in other pLGICs (see below), metal-ion modulation of DeCLIC may involve multiple allosteric sites.

At acidic pH without Ca^2+^, the open state shared the grid not with the resting-closed state, but with a seemingly heterogeneous class representing multiple configurations of the NTD (Figure 5B). Reconstruction of this partly disordered class [30] revealed contributors with individual NTD lobes asymmetrically displaced, generally outward from the pore axis, with visible similarity to models captured by MD. This alternative state was also consistent with SANS data, which provide structural information to lower resolution than cryo-EM, but without freezing on carbon-metal grids. Ensemble optimization of MD frames to fit SANS curvesindicated a substantial contribution by an extended-NTD conformation under all conditions, increasing at acidic pH upon Ca^2+^ depletion. Previous MD and SANS data [17] also indicate that alternative, largely extended-NTD conformations are present at neutral pH and in the presence of Ca^2+^, though their flexibility may render them difficult to resolve by cryo-EM or X-ray methods. The partly disordered structure captured here also features a relative clockwise rotation of the ECD and a closed pore, further indicating that NTD heterogeneity is anti-correlated with activation. A plausible model is that NTD disordering favors closed channels, whether by disruption of direct Ca^2+^ interactions or allosteric effects via the ECD. Calcium can partly stabilize the NTD, though acidification is needed to produce a prominent open population in cryo-EM or SANS experiments. Conversely, acidification in the absence of Ca^2+^ stabilizes some open channels, but also a substantial population of closed channels with disordered NTDs.

It remains unclear whether Ca^2+^ release is physiologically relevant to DeCLIC activity, or primarily a consequence of channel gating. Given that similar metal-ion sites are observed in several eukaryotic channels [41–45], elucidating this point may be of interest to understanding modulatory mechanisms across the pLGIC family. Interestingly, several of these receptors are also sensitive to pH [8–10], which could further influence metal-ion binding. The ECD-ion site in DeCLIC contributes to a prospective nonlinear permeation pathway, recently characterized also in eukaryotic and other prokaryotic pLGICs [28, 46–50]. In addition to mediating prospective Ca^2+^ block, this site contributes to an allosteric interface that undergoes substantial rearrangement upon Ca^2+^ depletion. Although conformational changes between the β1–β2 loop and loop F appear to be particularly dramatic in bacterial channels such as DeCLIC and sTeLIC [22], the equivalent region has been implicated in transducing ligand activation to pore opening across the pLGIC family [14]. It is plausible that transient binding of metal ions in the DeCLIC ECD modulates hyper-excitability in its native environment, analogous to desensitization in higher-organism pLGICs. Furthermore, pH variations in the salt-marsh environment, and/or in relation to symbiotic metabolism, could modulate affinity of metal-binding sites [51].

Despite the prokaryotic origins of DeCLIC, defining its gating and modulation mechanisms has proven complex. Apparent differences in open-state accessibility in cryo-EM, SANS, MD, and electrophysiology experiments highlight potentially critical influences of experimental conditions on the conformational landscape. For instance, solution-phase SANS profiles in several conditions are better fit by models with more heterogeneous, expanded NTDs than the relatively ordered states resolved by cryo-EM; this discrepancy illustrates intrinsic barriers to characterizing flexible modulatory domains. Although our open structure is evidently capable of ion permeation, MD simulations indicated a loss of charge selectivity at lower pH; this unanticipated result is likely overestimated by fixed protonation states and the uniform pH on external and internal sides of the membrane, both potential issues in modeling proton gating. Due in part to experimental challenges, we have observed substantial currents only in the absence of Ca^2+^, both in oocytes [16] and in lipid bilayers; it remains unclear how pH and Ca^2+^ availability in these configurations compare to structural biology experiments. Indeed, the present open state has only been identified at low pH, yet DeCLIC could be transiently activated at neutral pH in functional recordings [16], and SANS curves indicated a subpopulation of open channels under both pH conditions. These results echo recent reports of other pLGICs occupying different states in different membrane mimetics [37, 52], though at least in the case of DeCLIC, solubilization in lipid nanodiscs does not seem to facilitate opening under cryo-EM conditions [29].

Although our structure-function data do not provide direct insights into native DeCLIC function, it is interesting to speculate whether NTD ordering might be facilitated by periplasmic interaction partners. It is plausible that stable NTD interactions would promote function, given that channel opening is associated with a relatively stable NTD, and NTD truncation has been shown to slow gating [16]. Whatever its physiological role, the DeCLIC NTD offers a compelling case study in differential interactions of an accessory domain with distinct functional states of a pLGIC. The NTD adopts distinct conformations in the closed, open, and partly disordered structures reported in this work, recapitulating extra-membrane domain contractions that have been consistently implicated in pLGIC gating [14]. Our data further illustrate the power of biophysical methods including cryo-EM, electrophysiology, MD, and SANS to elucidate complex motions of the DeCLIC NTD. Computational and NMR approaches have similarly proved instrumental in characterizing the ICD of eukaryotic channels [53], which appears to be similarly flexible and modulatory.

## Materials and Methods

### DeCLIC Expression and Purification

DeCLIC was expressed and purified as in [17]. To summarize, DeCLIC-MBP in pET-20b was transformed into C43(DE3) *E. coli* and cultured overnight at 37° C. Overnight culture was inoculated 1:100 into 2xYT media with 100 µg/mL ampicillin. The culture was grown at 37° C until it reached OD_600_ of 0.8, at which point it was induced with 100 µM isopropyl-β-D-1-thiogalactopyranoside (IPTG). The temperature was lowered to 25° C and the cells were left shaking overnight. Once harvested, cells were resuspended in buffer A (300 mM NaCl, 20 mM Tris-HCl pH 7.4) supplemented with protease inhibitors, 20 µg/mL DNase I, 1 mg/mL lysozyme and 5 mM MgCl_2_. After sonication the membranes were harvested by ultracentrifugation, followed by solubilization in 2% n-dodecyl-β-D-maltoside (DDM). Fusion proteins were purified by amylose affinity (NEB), eluting in buffer B (buffer A with 0.02% DDM) and eluted with 2–20 mM maltose. Size exclusion chromatography (SEC) in buffer B was used to further separate the fusion protein from endogenous maltose binding protein. After overnight thrombin digestion, DeCLIC was isolated from its fusion partner with a final size exclusion run in Buffer C (300 mM NaCl, 20 mM Citrate pH 5, 0.02% DDM) either with 10 mM Ca^2+^ or 10 mM EDTA. Finally, the protein was concentrated to 3–5 mg/mL by centrifugation.

### Cryo-EM Sample Preparation and Data Acquisition

The same freezing protocol was applied for both of the conditions. 3 µl of DeCLIC sample was applied to a glow-discharged quantifoil 1.2/1.3 Cu 300 mesh grid (Quantifoil Micro Tools), followed by 30 s waiting time at 4° C and 100% humidity. Sample was then blotted for 3 s and plunge-frozen into liquid ethane using a FEI Vitrobot Mark IV. Micrographs were collected on an FEI Titan Krios 300 kV microscope with a Gatan K3-Summit direct detector camera. Movies were collected at nominal 105,000x magnification, equivalent to a pixel spacing of 0.8617Å. A total dose of 42 e^-^/Å^2^ was used to collect 40 frames over 2 sec, using a nominal defocus range covering −1.6 to −3.0 µm (for the no Ca^2+^ dataset) and −1.4 to −3.0 µm (for the Ca^2+^ present dataset) in steps of 0.2 µm.

### Image Processing

Data processing was performed through the Relion 4.0-beta-2 pipeline [54]. Motion correction was performed with Relion’s own implementation [55]. Defocus was estimated from the motion corrected micrographs using CtfFind4 [56]. References for autopicking were generated with a 2D classification performed on a subset of manually picked particles. Following the autopicking on a full dataset the particles were extracted and binned 4 times before the initial reference was generated. In silico purification was performed through multiple rounds of 2D- and 3D-classification as well as 3D auto-refinement. Per-particle CTF parameters were estimated from the resulting reconstruction using Relion 4.0-beta-2. Global beam-tilt was estimated from the micrographs and correction applied. Following particle polishing a final 3D auto-refinement was performed using a soft mask. Detergent micelle density was subtracted with a mask generated from a DeCLIC structure (PDB ID: 7Q3H, [17]), followed by post-processing with the same mask. Local resolution was estimated using the Relion implementation.

### Model Building

Cryo-EM structures in the presence and absence of Ca^2+^ were built starting from a monomer of previously solved DeCLIC cryo-EM structure (PDB ID: 7Q3H, chain A [17]), where the missing residues in the NTD region were added with Modeller [57]. To refine the models, Phenix 1.21rc1-5109 [58] real-space refinement was used. 5-fold symmetry was imposed through NCS restraints, which were detected from the reconstructed cryo-EM maps. Models were manually adjusted in Coot 0.9.5 EL [59] and re-refined in Phenix ([58] until the model statistics reached satisfactory values (Supplementary Table **??**). Atoms were removed where the density did not allow for confident building (Supplementary Table **??**). Buried surface area around NTD lobes was calculated for each subunit using the “measure buriedarea” command in ChimeraX [60], using either the residues of NTD1 and NTD2, or NTD2 and EDC.

### Heterogeneous Reconstruction

Partly disordered particles from the pH-5 EDTA data set were selected based on their 2D classes (Figure **??** A) and used to train an 8D latent variable model for 50 epochs with a 1024 x 3 encoder and decoder architecture in cryoDRGN 2.3 ([30]. Next, the default 20 *k*-means cluster center densities produced with the cryoDRGN analysis tool were manually clustered into 4 groups based on structural similarities, and the particles belonging to the *k*-means clusters of these 4 groups were extracted and then refined in Relion using 3D-auto refine (Figure **??** B). Additionally, given the evident heterogeneity in this set of particles revealed by cryoDRGN analysis, the 2D classes in **??**A were divided into two major classes that were both refined by multiple rounds of 3D auto-refine and 3D classification, resulting in two low-resolution (7 Å) reconstructions with clearly different NTD arrangements (Figure **??**C). Applying C5 symmetry resulted in slightly higher resolution densities (5 Å) that were superimposable in the ECD and TMD, only differing in the NTD.

### Lipid Bilayer Recording

Lipid bilayer recordings were carried out on the Orbit mini instrument (Nanion Technologies GmbH) on a MECA4 recording chip. For all experiments, Ringer’s buffer at pH 5 or pH 7, with or without 100 mM CaCl_2_, was applied to the chip before the formation of a bilayer using 1,2-diphytanoyl-sn-glycero-3-phosphocholine (DPhyPC) phospholipids (Avanti Polar Lipids Inc.) dissolved in octane to a concentration of 10 mg/ml. Upon the formation of stable bilayers (capacitance above 7 pF) in all four channels, voltage was clamped at +50 mV, and purified DeCLIC was added to a final concentration of 6 ng/ml. Electrophysiology traces were recorded with Elements Data Reader (Elements SRL) for at least 5 minutes. Each experiment was repeated 10 times, and the data from bilayers that retained good quality throughout the recording (19–21 bilayers per condition, defined as them breaking from a zap) was analysed in Clampfit (Axon, Molecular Devices). A 50x data reduction was applied to each trace (substitute average), resulting in a sampling rate of one frame per 40.96 ms. Then, a so-called event detection threshold search was performed to identify instances of activity in all traces, where a deviation of 0.2 pA from the baseline (minimum conductance 4 pS) for at least 165 ms triggered the start of an event, and a return to less than 0.2 pA from the baseline for at least 165 ms was defined as the end of an event. Fractional activity per recorded bilayer, event counts and duration were quantified and plotted using Prism 10.2.3 (GraphPad) and are summarized in **??**. Significance was determined by two-tailed unpaired *t*-test with Welch’s correction.

### Molecular Dynamics Simulations

MD simulations were launched from cryo-EM structures reported in this work, using full-length models prior to removal of residues with insufficient density. For the closed structure, five bound Ca^2+^ ions were retained. Local pK_a_ was estimated for both structures using Propka3 [31, 32]; based on these estimated values, protonation states most likely at either pH 5 (referred to as protonated) or pH 7 (deprotonated) were applied to generate the corresponding systems. Simulation systems were set up using the Charmm-GUI membrane builder [61, 62], embedding the protein in a POPC bilayer, adding sodium and chloride ions to neutralize the system and approximate 150 mM concentration, and solvating in TIP3 water. Hydrogen mass re-partitioning [63] and the Charmm36m force-field were used [64], and simulations were performed in Gromacs v. 2021.3 [65]. Systems were energy-minimized, followed by equilibration with gradual release of restraints. Four replicate production simulations of 1000 ns were performed for each system.

Root-mean-square deviations (RMSDs) were calculated using Gromacs, for C-alpha atoms of full subunits and for individual structural regions, using two different alignments: the selection itself, or the ECD-TMD region. Distances and center-of-mass positions were tracked using Vmd [66]. For tracking ions in the pore, an initial identification of relevant ions was made using a Vmd script to report the atom indices of ions within a cut-off distance of the pore axis which passed the z-value of one of the pore constrictions. Spuriously reported ions were identified by visual inspection, and the z-coordinates of the remaining selection of ions were tracked using Vmd. Pore profiles for the simulations and for the structures were calculated using Chap [67]. Vmd [66] and UCSF ChimeraX [60] were used to render images.

### Small-Angle Neutron Scattering

The SEC-SANS experiments [68] were performed using a paused-flow approach [34] at the D22 beamline of Institute Laue–Langevin. The experimental procedure was as previously described [17], using gel filtration in-line with SANS measurements [69, 70] and match-out deuterated DDM (d-DDM) [33]. The detector distances for the multidetector banks were 1.4 m and 8 m, covering a Q-range of 0.0065 – 0.73 Å^-1^. The definition of Q used was Q=(4π/λ)sin(*θ*), where 2*θ* is the scattering angle and λ is the wavelength. Two buffer conditions were used: with Ca^2+^ (D_2_O, 150 mM NaCl, 20 mM Citrate*·*HCl, 10 mM CaCl_2_, 0.5 mM d-DDM), and without Ca^2+^ (D_2_O, 150 mM NaCl, 20 mM Citrate*·*HCl, 10 mM EDTA, 0.5 mM d-DDM). Further sample details and collection parameters are available in Supplementary Tables **??** and **??**. Data reduction was performed using Grasp version 10.02 [71, 72]. Data collected prior to the protein peak provided the scattering used for buffer subtraction. Concentration normalization was done using the average protein concentration during the measurement, which was calculated from the co-recorded chromatogram and the extinction coefficient provided by ProtParam [73] from the amino acid sequence. Additionally a small constant was subtracted to adjust the background.

Guinier analysis was used to calculate radius of gyration (R_g_) (Supplementary Figure **??**, Supplementary Table **??**). Pairwise distance distributions were calculated for experimental data using BayesApp [74], and for all-atom models using CaPP (Calculating Pair distance distribution functions for Proteins) [75]. Pepsi-SANS [76] was used to calculate theoretical scattering curves and their fits to the experimental scattering profiles. Linear combinations of theoretical scattering curves were calculated in Python3. Ensemble optimization was performed using Gajoe from Eom [35, 36], providing theoretical scattering intensities calculated with Pepsi-SANS as the model pool. The ensemble size was set to 28 (approximately 1/100th the size of the model pool), repeats of models were allowed in the ensemble, constant subtraction was disabled, remaining settings were kept at default values. Gajoe was run five times per fitted SANS dataset. A summary of equations and software used is available in Supplementary Table **??**.

## Data Availability

Cryo-EM density maps of the pentameric ligand-gated ion channel DeCLIC in detergent micelle have been deposited in the Electron Microscopy Data Bank under accession numbers EMD-50745 (open, Ca^2+^), EMD-50744 (closed, Ca^2+^), EMD-50746 (open, EDTA) and EMD-50747 (partly disordered, EDTA). Each deposition includes the cryo-EM unsharpened maps, both half-maps and the mask used for final FSC calculation. Coordinates of the models have been deposited in the Protein Data Bank, with the accession numbers 9FTH (open, Ca^2+^), 9FTG (closed, Ca^2+^), 9FTI (open, EDTA) and 9FTJ (partly disordered, EDTA). The raw experimental SANS data is available from https://doi.ill.fr/10.5291/ILL-DATA.EASY-861, and processed SANS data are available in the Small Angle Scattering Biological Data Bank [77] (SASBDB) as entries SASDVX3 (with Ca^2+^, https://www.sasbdb.org/data/SASDVX3) and SASDVY3 (with EDTA, https://www.sasbdb.org/data/SASDVY3). MD simulation trajectories and alternative-NTD conformation cryo-EM densities are available at Zenodo.org at https://doi.org/10.5281/zenodo.11383747. Previously published data utilized in this work are available at SASBDB (SASDNG5 and SASDNH5) and Zenodo (https://doi.org/10.5281/zenodo.6022369).

## Acknowledgments

The authors would like to thank the Swedish Cryo-EM National Facility staff, especially Marta Carroni and Stefan Fleischmann from Stockholm, for kind assistance with data collection. The authors would also like to acknowledge the Institut Laue-Langevin for development of SEC-SANS and allocated beam time at the D22 instrument, and in particular Anne Martel and Lionel Porcar for assistance with data collection. This work was supported by grants from SwedNess, the Knut and Alice Wallenberg Foundation, the Swedish Research Council, the Swedish e-Science Research Centre, and the BioExcel Center of Excellence. Cryo-EM data were collected at the Swedish national cryo-EM facility funded by the Knut and Alice Wallenberg Foundation, Erling Persson and Kempe Foundations. Computational resources were provided by the Swedish National Infrastructure for Computing (SNIC).

## Author Contributions

Conceptualization: UR, OA, ML, MD, RJH, EL; methodology: UR, OA, ML, RJH; software: UR, OA, ML; validation: UR, OA, ML, RJH; formal analysis: UR, OA, ML, RJH; investigation: UR, OA, ML; resources: UR, OA, ML, RJH; data curation: UR, OA, ML, RJH; original draft: UR, ML; review and editing: UR, OA, ML, MD, RJH, EL; visualization: UR, OA, ML; supervision: RJH, EL; project administration: RJH; funding acquisition: EL.

## Competing intrests

The authors declare that they have no conflict of interest.

