## Supplementary Information for "Structure and dynamics of a multidomain ligand-gated ion channel revealed under acidic conditions"

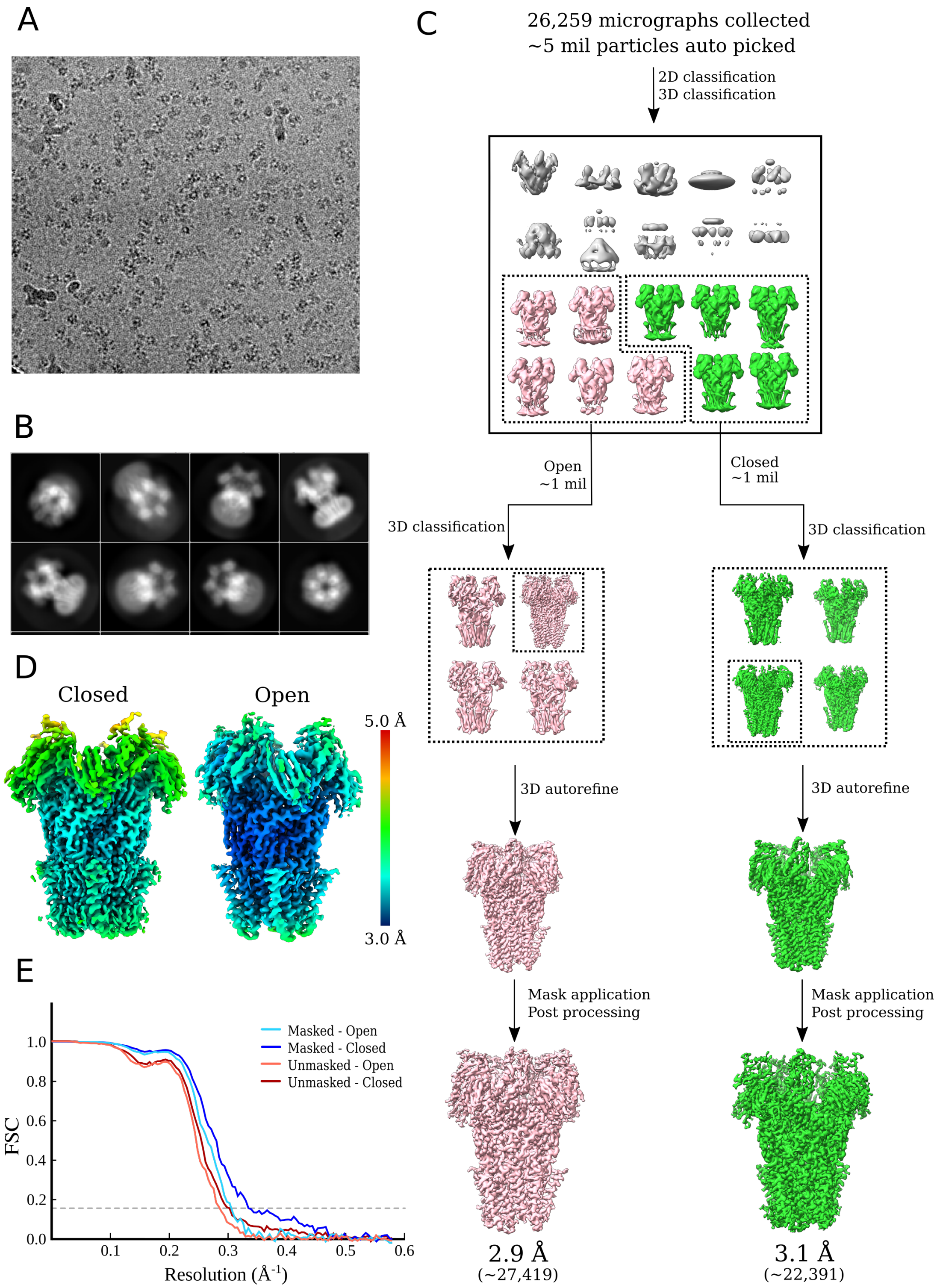


**Figure S1: Cryo-EM data processing overview for DeCLIC at pH 5 with 10 mM calcium.** (A) Representative micrograph. (B) Representative good 2D classes. (C) Graphical overview of the cryo-EM processing pipeline. (D) Electron densities colored by local resolution. (E) Fourier shell correlation (FSC) curves for masked and unmasked densities assigned to open and closed states.

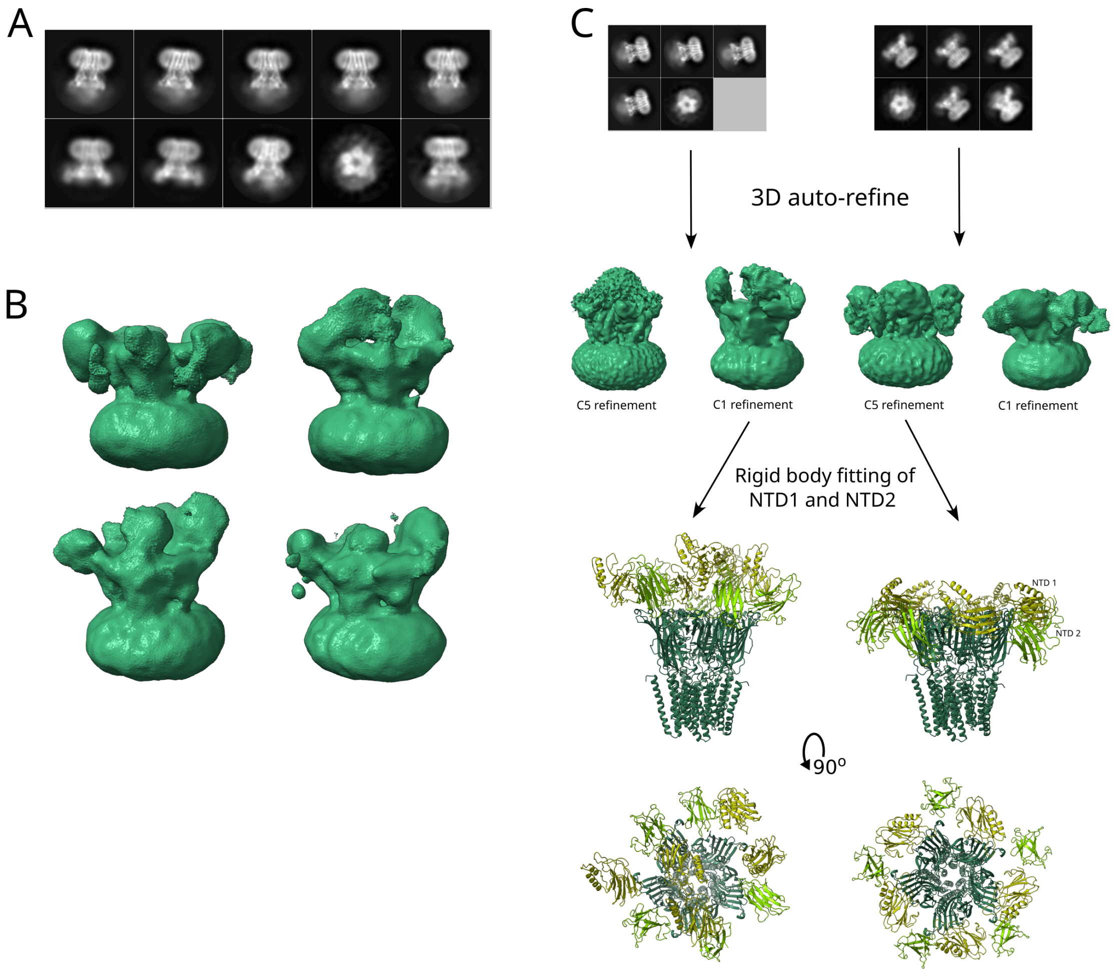


**Figure S2: Heterogeneity in the partly disordered class of DeCLIC resolved at pH 5 without calcium.** (A) 2D classes of partly disordered particles used to train a latent variable model in cryoDRGN [30]. (B) Densities produced by 3D-auto refine in RELION using particles selected based on the cryoDRGN latent space encoding and clustering, showing diverse arrangements of the NTD. (C) Pipeline for refinement of the two major disordered clusters and rigid body fitting of the NTD lobes into the resulting densities, showing two distinct conformations of the NTD.

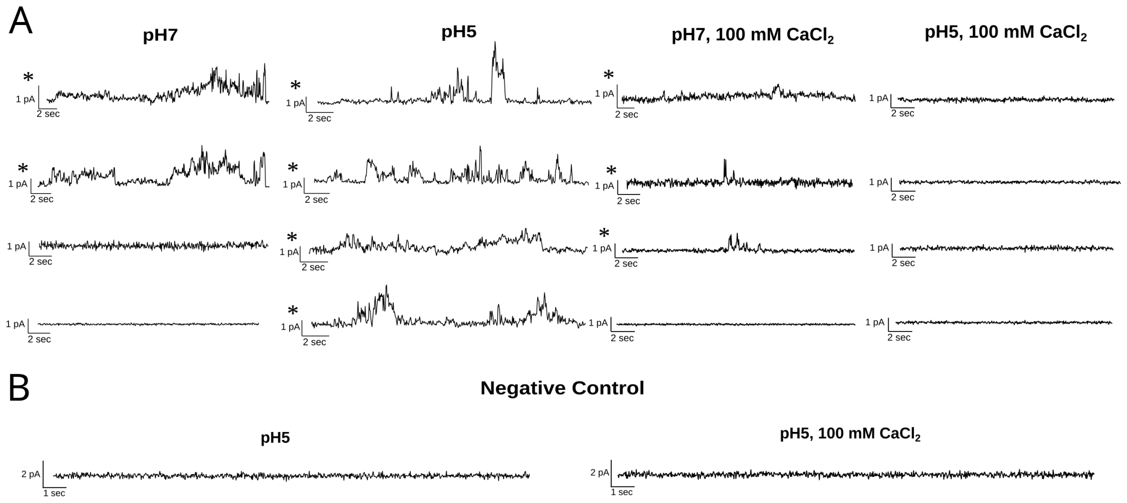


**Figure S3: Lipid bilayer recording sample traces.** (A) Sample traces recorded from DeCLIC at pH 5 and 7 with and without 100 mM $CaCl_2$. Traces from membranes with detected activity are marked with an asterisk (*). Traces include samples from all membranes with detected activity not shown in Figure 2, with the exception of pH 5 without calcium, where an additional three membranes showed evidence of activity. (B) Representative sample traces from recordings of membranes without DeCLIC added, at pH 5 with (right) and without (left) 100 mM $CaCl_2$.

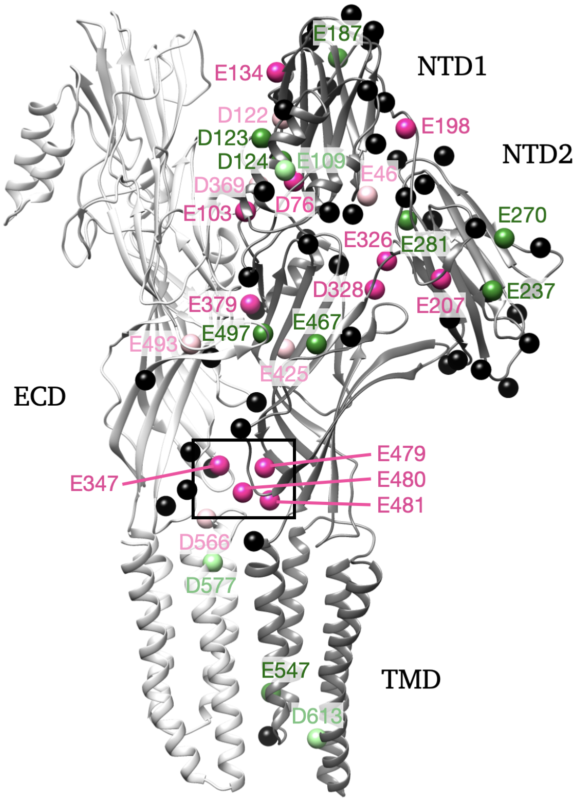


**Figure S4: Acidic residues at a single DeCLIC subunit interface.** Primary and complementary DeCLIC subunits are depicted as light and dark gray ribbons respectively, viewed from the membrane plane; for clarity, the distal three subunits are hidden. Spheres represent Cβatoms of all 76 Asp and Glu residues associated with a single interface. Of these, 30 residues predicted by PropKa to protonate at pH 5 in at least one state are labeled. Magenta labels indicate the local pKa increases by at least 0.5 units in the open versus closed states, consistent with a role in proton activation. Green labels indicate the local pKa increases at least 0.5 units in closed versus open states, an inverse correlation with proton activation. Light pink and light green labels indicate relative pKa increases by less than 0.5 units in open and closed states, respectively. Boxed region corresponds to zoom region in Figure 2A at the subunit/domain interface.

| Position | Type | Closed State | | | Open State | | | Region |
|---|---|---|---|---|---|---|---|---|
| | | pKa | Protonation pH 7 | Protonation pH 5 | pKa | Protonation pH 7 | Protonation pH 5 | |
| 35 | Glu | 4.64 | | | 4.76 | | | |
| 46 | Glu | 4.96 | | | 5.32 | | x | |
| 47 | Asp | 4.44 | | | 4.15 | | | |
| 57 | Asp | 3.97 | | | 4.28 | | | |
| 62 | Asp | 4.02 | | | 4.36 | | | |
| 76 | Asp | 5.83 | | x | 8.28 | x | x | |
| 84 | Glu | 3.97 | | | 4.65 | | | |
| 85 | Glu | 4.68 | | | 4.23 | | | |
| 87 | Asp | 3.52 | | | 4.27 | | | |
| 97 | Glu | 3.46 | | | 4.68 | | | |
| 103 | Glu | 4.6 | | | 7.49 | x | x | |
| 105 | Asp | 4.62 | | | 4.02 | | | |
| 109 | Glu | 5.11 | | x | 4.95 | | | NTD1 |
| 115 | Asp | 3.85 | | | 3.32 | | | |
| 122 | Asp | 5.77 | | x | 6.10 | | x | |
| 123 | Asp | 5.09 | | x | 3.53 | | | |
| 124 | Asp | 7.37 | x | x | 6.48 | | x | |
| 134 | Glu | 4.70 | | | 5.48 | | x | |
| 140 | Asp | 4.25 | | | 3.88 | | | |
| 157 | Asp | 2.24 | | | 3.68 | | | |
| 183 | Glu | 4.62 | | | 4.65 | | | |
| 185 | Glu | 4.76 | | | 4.9 | | | |
| 187 | Glu | 5.17 | | x | 4.44 | | | |
| 189 | Glu | 4.38 | | | 4.33 | | | |
| 198 | Glu | 4.47 | | | 5.24 | | x | |
| 202 | Glu | 4.24 | | | 4.80 | | | |
| 207 | Glu | 4.22 | | | 6.34 | | x | |
| 209 | Glu | 4.81 | | | 4.49 | | | |
| 213 | Asp | 2.74 | | | 3.24 | | | |
| 215 | Asp | 3.78 | | | 4.40 | | | |
| 230 | Asp | 4.48 | | | 4.30 | | | |
| 237 | Glu | 5.38 | | x | 4.69 | | | |
| 241 | Glu | 4.71 | | | 4.58 | | | |
| 251 | Asp | 4.89 | | | 4.83 | | | |
| 263 | Asp | 3.55 | | | 3.29 | | | NTD2 |
| 267 | Glu | 4.94 | | | 4.64 | | | |
| 270 | Glu | 5.38 | | x | 4.51 | | | |
| 276 | Asp | 4.16 | | | 4.39 | | | |
| 278 | Asp | 4.00 | | | 4.05 | | | |
| 281 | Glu | 5.14 | | x | 3.56 | | | |
| 287 | Asp | 5.05 | | | 4.76 | | | |
| 290 | Asp | 3.31 | | | 3.85 | | | |
| 294 | Asp | 3.17 | | | 3.69 | | | |
| 297 | Glu | 4.65 | | | 4.53 | | | |
| 313 | Glu | 4.40 | | | 4.69 | | | |

**Figure S5: Predicted protonation behavior of acidic residues in the DeCLIC NTD.** Local pKa's and predicted protonation states at pH 7 and pH 5 for Asp and Glu residues in closed and open structures as calculated in PropKa [32].

| Position | Type | Closed State pKa | Closed State Protonation pH 7 | Closed State Protonation pH 5 | Open State pKa | Open State Protonation pH 7 | Open State Protonation pH 5 | Region |
|---|---|---|---|---|---|---|---|---|
| 326 | Glu | 4.73 | | | 5.99 | | x | |
| 328 | Asp | 5.47 | | x | 6.28 | | x | |
| 337 | Asp | 4.86 | | | 4.64 | | | |
| 343 | Asp | 3.73 | | | 3.83 | | | |
| 347 | Glu | 6.12 | | x | 6.76 | | x | $Ca^{2+}$ site |
| 361 | Asp | 4.31 | | | 4.33 | | | |
| 369 | Asp | 5.85 | | x | 6.07 | | x | |
| 379 | Glu | 5.29 | | x | 6.19 | | x | |
| 380 | Asp | 4.26 | | | 5.00 | | | |
| 389 | Glu | 4.76 | | | 4.36 | | | |
| 418 | Asp | 3.97 | | | 4.16 | | | |
| 425 | Glu | 4.94 | | | 5.18 | | x | ECD |
| 435 | Asp | 3.91 | | | 3.97 | | | |
| 437 | Asp | 4.45 | | | 3.93 | | | |
| 444 | Asp | 4.04 | | | 3.65 | | | |
| 453 | Asp | 4.41 | | | 4.67 | | | |
| 465 | Glu | 4.72 | | | 4.75 | | | |
| 467 | Glu | 5.11 | | x | 3.94 | | | |
| 475 | Asp | 3.99 | | | 3.81 | | | |
| 479 | Glu | 5.14 | | x | 7.21 | x | x | |
| 480 | Glu | 5.61 | | x | 6.94 | | x | $Ca^{2+}$ site |
| 481 | Glu | 5.09 | | x | 5.88 | | x | |
| 490 | Glu | 4.70 | | | 4.68 | | | |
| 493 | Glu | 4.97 | | | 5.28 | | x | |
| 497 | Glu | 6.05 | | x | 5.17 | | x | |
| 516 | Glu | 4.61 | | | 4.48 | | | |
| 541 | Asp | 3.38 | | | 3.82 | | | |
| 547 | Glu | 6.19 | | x | 4.892 | | | TMD |
| 566 | Asp | 4.65 | | | 5.04 | | x | |
| 577 | Asp | 5.03 | | x | 4.70 | | | |
| 613 | Asp | 5.38 | | x | 5.22 | | x | |

**Figure S6: Predicted protonation behavior of acidic residues in the DeCLIC ECD and TMD.** Local pKa's and predicted protonation states at pH 7 and pH 5 for Asp and Glu residues in closed and open structures as calculated in PropKa.

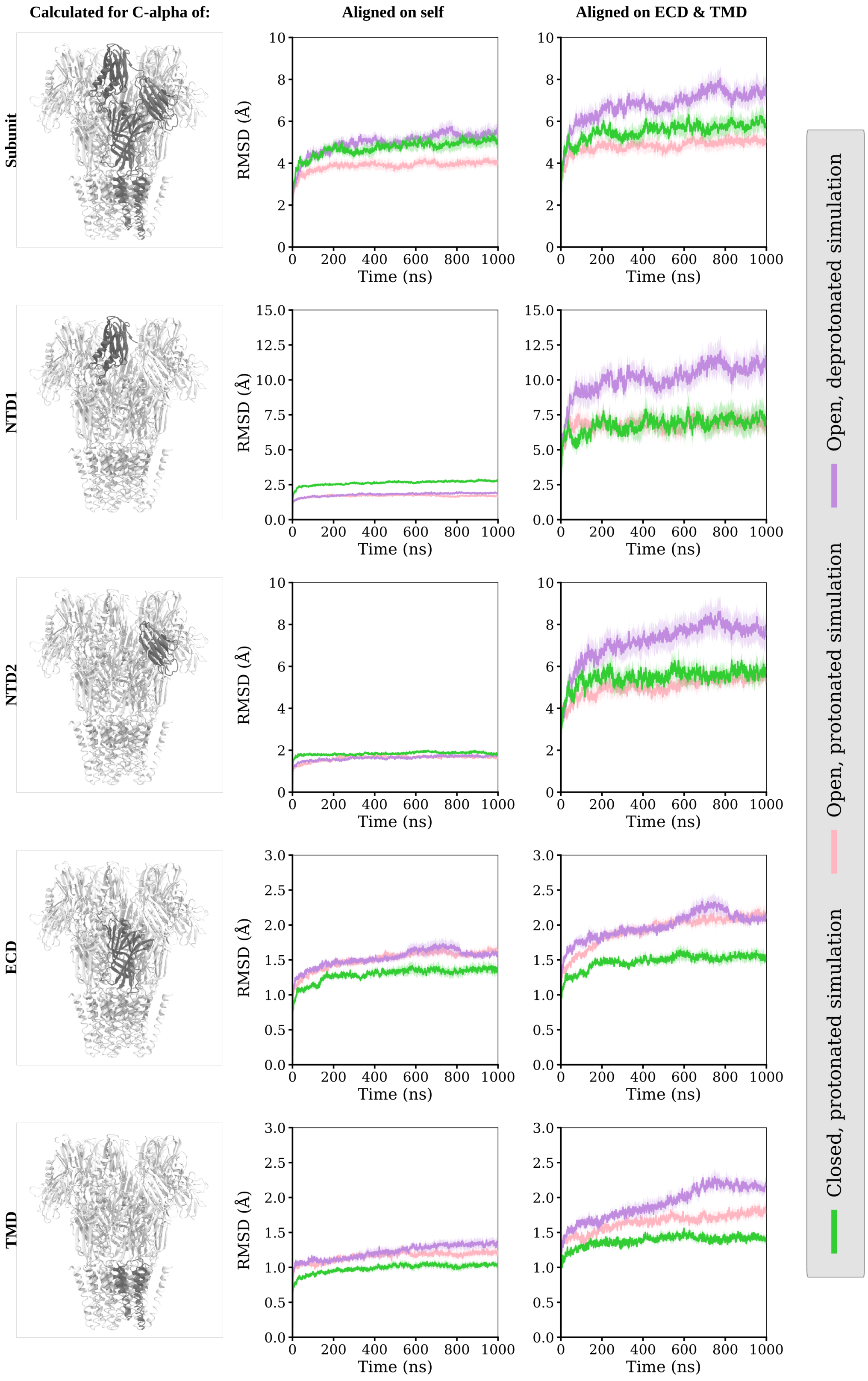


**Figure S7: Overall and regional stability in DeCLIC simulations.** *Left,* DeCLIC models with the region used for alignment in each row highlighted black. *Center,* RMSD of C-alpha atoms in an entire DeCLIC subunit (top row), an isolated NTD1 (second row) or NTD2 lobe (third row), or a single-subunit ECD (fourth row) or TMD (bottom row), based on models aligned on the indicated region. Solid lines represent mean values over all five subunits across four replicates; shading indicates standard error of the mean. Systems include the closed structure at pH 5 (protonated, lime), open structure at pH 5 (protonated, pink), and open structure at pH 7 (deprotonated, purple). *Right,* RMSD plots as in center, based on models aligned on the ECD-TMD region of a given chain. Elevated RMSDs in both NTD lobes, when aligning on the ECD-TMD region versus the lobes themselves, indicate rigid-body translations relative to the minimal channel.

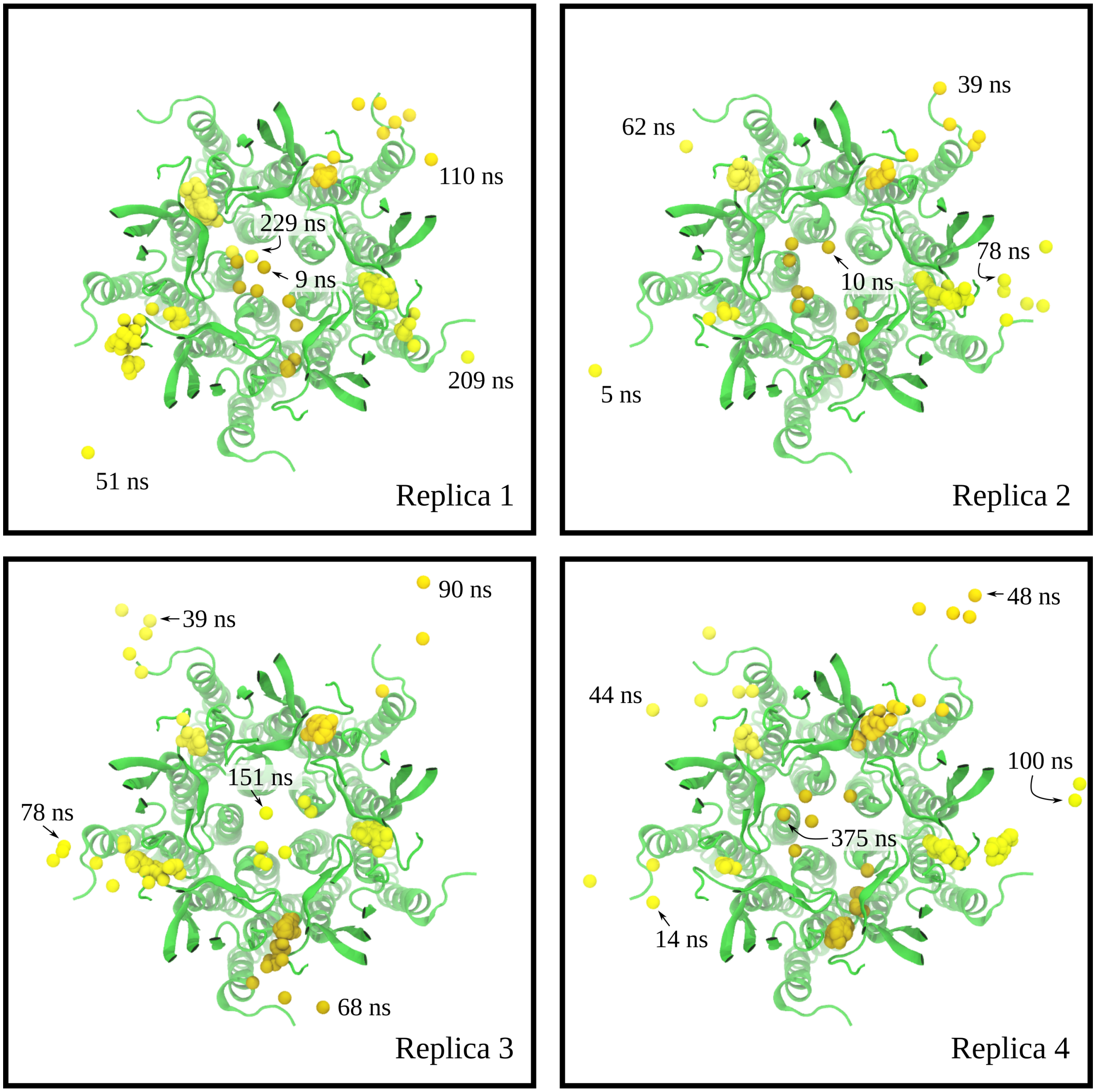


**Figure S8: Dissociation of calcium ions at low pH.** Calcium ions (yellow) in four replicate simulations of the closed structure determined at pH 5 with calcium, rendered as one sphere per ns from the start of the simulation until shortly after it dissociates from the protein. Labels indicate timestamps for the final render shown for each ion. DeCLIC is representated as a single initiating snapshot (green ribbons). Each system is viewed from the extracellular side, with the NTD and part of the ECD hidden for clarity.

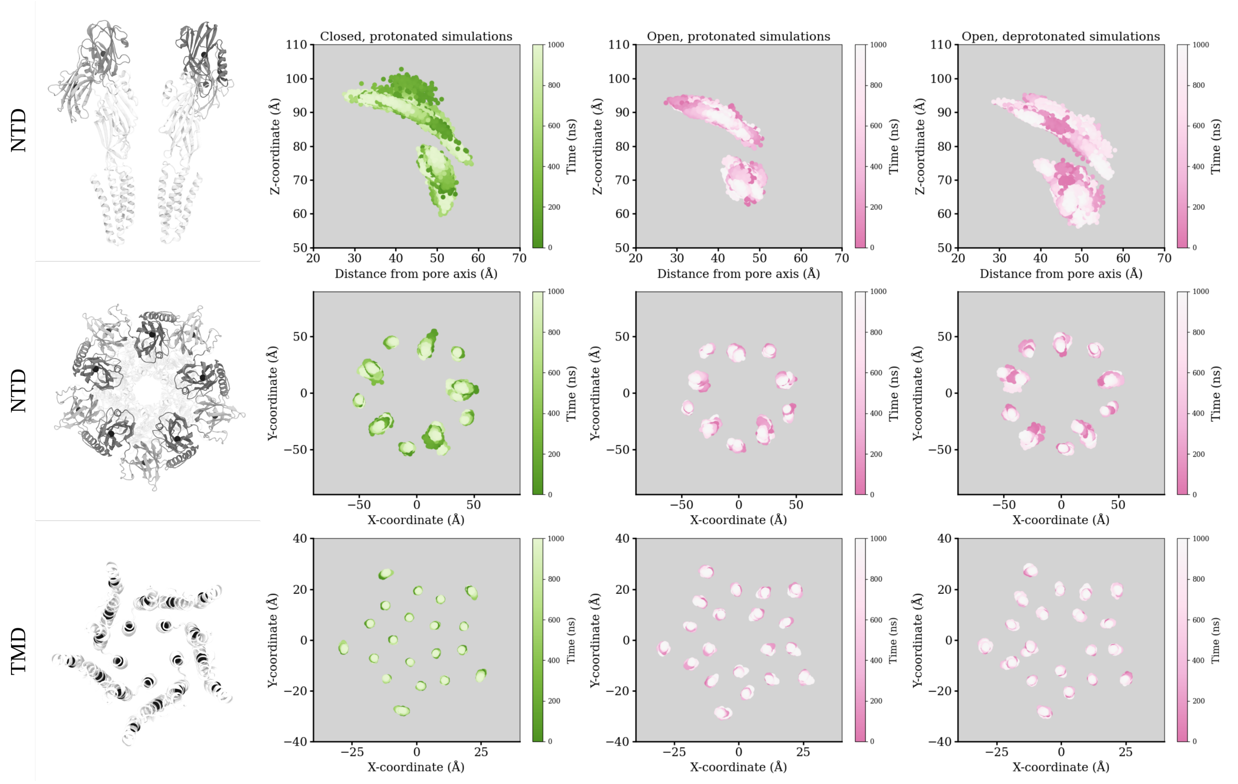


**Figure S9: NTD and TMD dynamics in DeCLIC simulations.** Top, center-of-mass positions of single-subunit NTDs, Renders illustrating the protein and tracked centers of mass are shown in the first column, with black spheres showing individual centers of mass of the selections. The NTD-lobes in the renders are highlighed in gray, and in the side view only two subunits are shown for clarity. The first row shows the distance from the pore axis and Z-coordinate for the NTD-lobes, the second row shows the X- and Y-coordinates of the NTD-lobes. The last row shows the X- and Y-coordinates for the transmembrane helices. The second column shows data from the protonated closed simulations (green), the third column for the protonated open simulations (pink), and the fourth column for the deprotonated open simulations (pink). Data points from all subunits and from four replicas per simulation condition are shown.

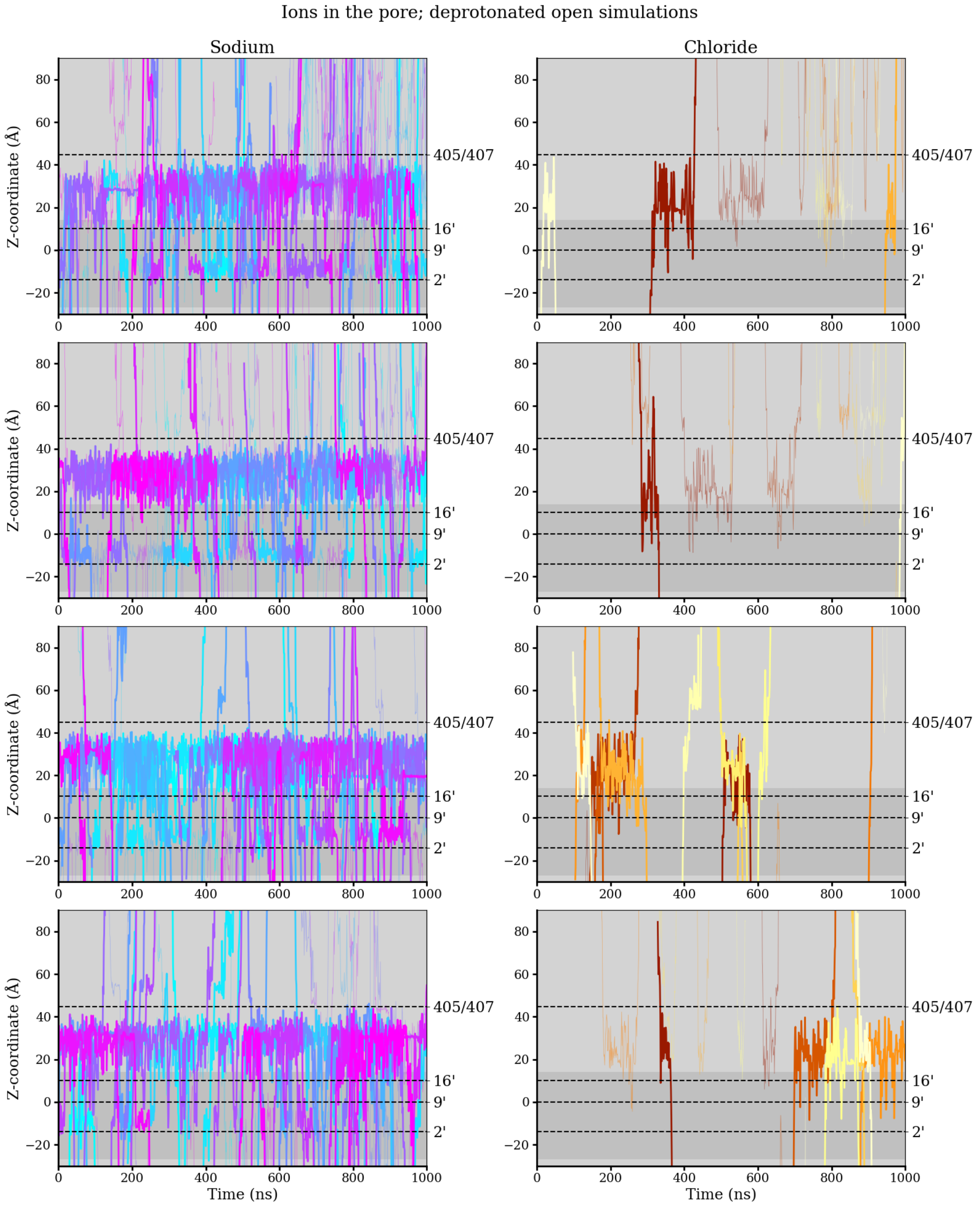


**Figure S10: Ion permeation in deprotonated simulations of the low pH open structure.** Z-coordinates as a function of time for ions inside the pore or extracellular lumen, the z-coordinate of 2’ defining the lowest point of the pore, and the constriction formed by residues 405 and 407 defining the upper point of the extracellular lumen. The z-range of the membrane is indicated by a darker gray background, thicker lines indicate ions which pass the 2’, 9’, and 16’ constrictions (dashed horizontal lines) in the transmembrane domain, thin lines indicate ions which enter the central conduction pathway but which do no fully transverse the transmembrane region. Each row represents data from one simulation replica, the left column the sodium ions in the pore, and the right column shows the chloride ions in the pore.

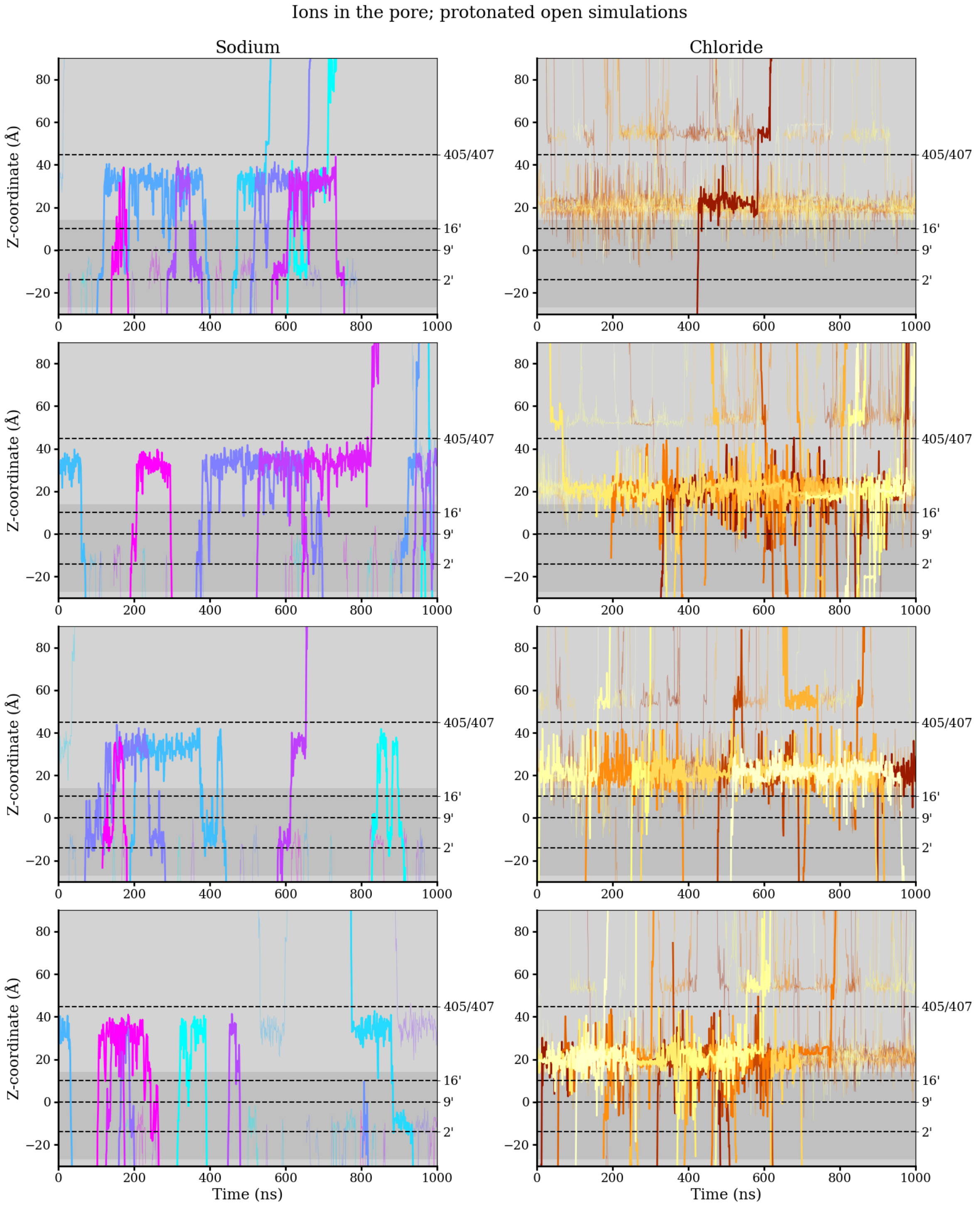


**Figure S11: Ion permeation in protonated simulations of the low pH open structure.** Z-coordinates as a function of time for ions inside the pore or extracellular lumen, the z-coordinate of 2' defining the lowest point of the pore, and the constriction formed by residues 405 and 407 defining the upper point of the extracellular lumen. The z-range of the membrane is indicated by a darker gray background, thicker lines indicate ions which pass the 2', 9', and 16' constrictions (dashed horizontal lines) in the transmembrane domain, thin lines indicate ions which enter the central conduction pathway but which do no fully transverse the transmembrane region. Each row represents data from one simulation replica, the left column the sodium ions in the pore, and the right column shows the chloride ions in the pore.

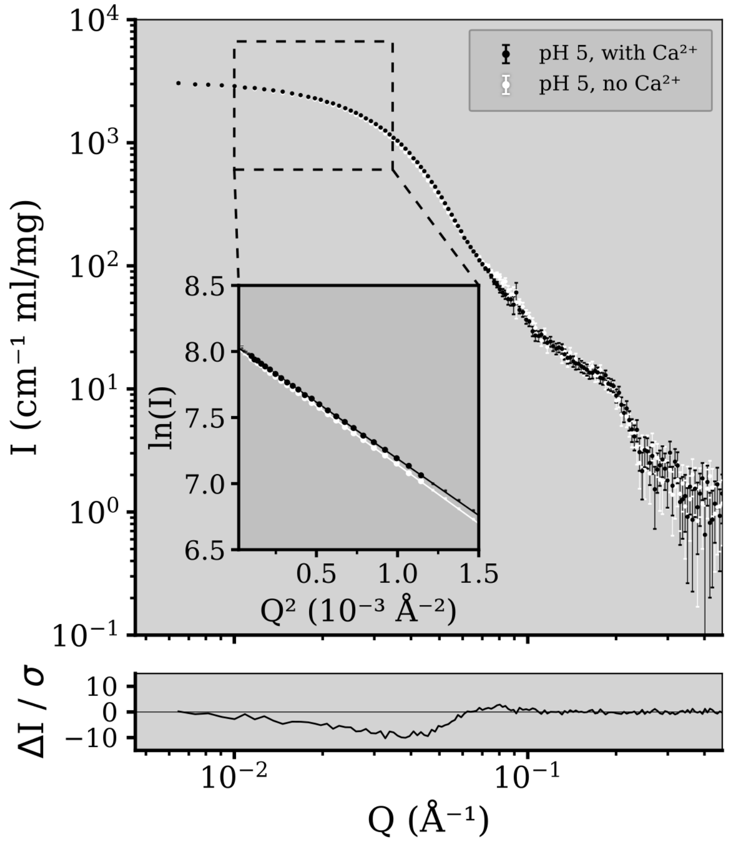


**Figure S12: Small-angle neutron scattering from DeCLIC at pH 5.** The main panel shows the scattering profiles from DeCLIC at pH 5 in the presence (black) and absence (white) of calcium. Dashed box indicates data points used for Guinier analysis, and the insert shows the Guinier plot; where experimental data-points are shown as dots (large dots were included in the Guinier fits), and the Guinier fits are shown as lines. The lower panel shows the error weighted residual between the two experimental curves.

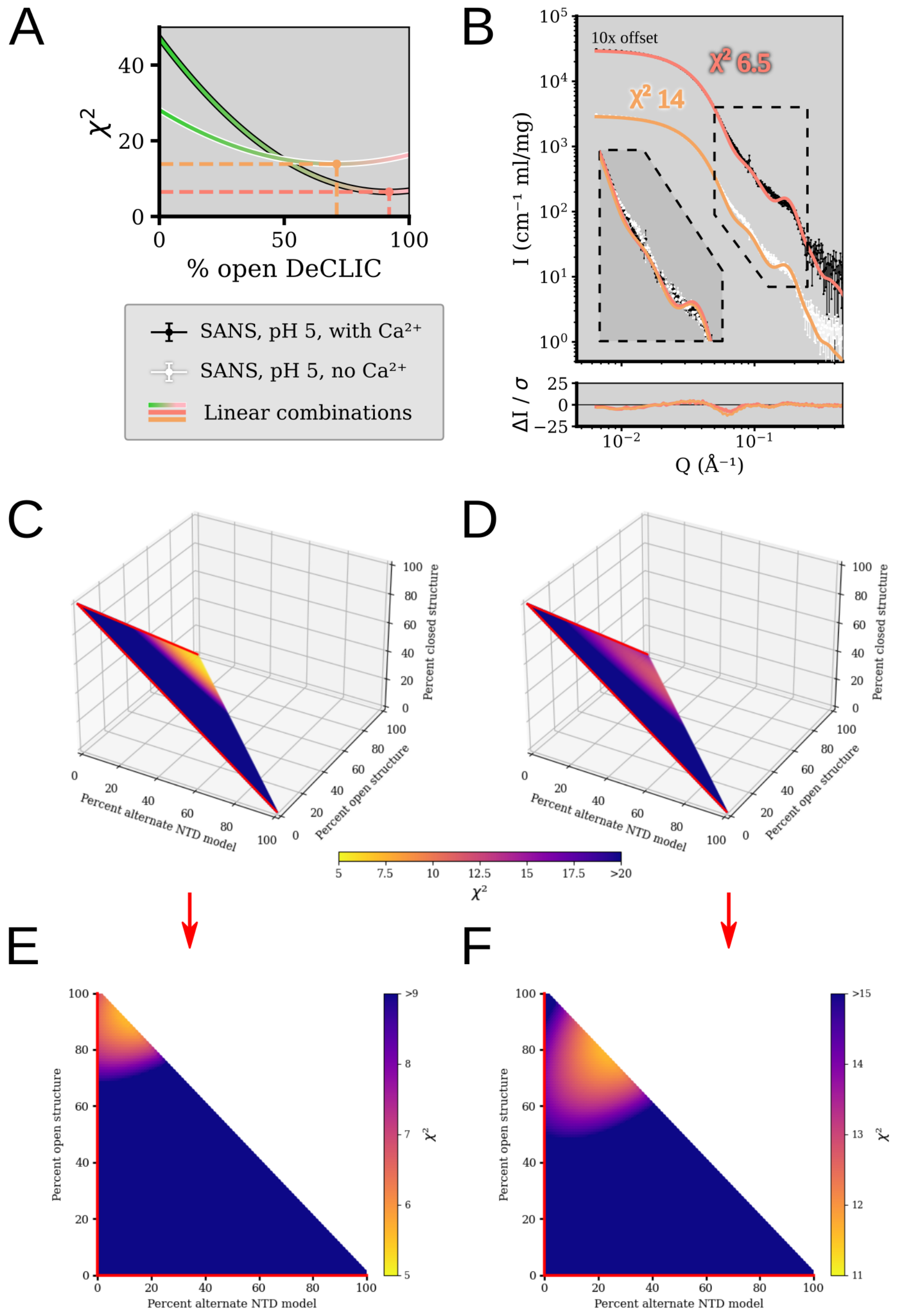


**Figure S13: Fits to SANS data for linear combination models from cryo-EM at pH 5.** (A) Goodness of fit as a function of percent open DeCLIC included in a linear combination of the pH 5 closed and open structures. Comparison to the with calcium data-set with black outline, and to the no calcium data-set with white outline. Dashed lines indicate the best combination for respective data-sets. (B) Experimental scattering profiles (with calcium dataset off-set by x10) together with the best linear combination of the pH 5 closed and open structures. The insert shows a zoom on the main feature, and the lower panel the error weighted residuals. (C) Reduced $\chi^2$ goodness of fit (colormap) as a function of varying the contribution of the open structure, closed structure, and the "up" alternate NTD model, in a linear combination fit to the SANS profile from DeCLIC at pH 5 in the presence of calcium, and (D) to the SANS profile from DeCLIC at pH 5 in the absence of calcium. (E) Projection of A on the plane of the contribution from the open and alternative NTD-conformation to the linear combination. (F) Projection like E, for D.

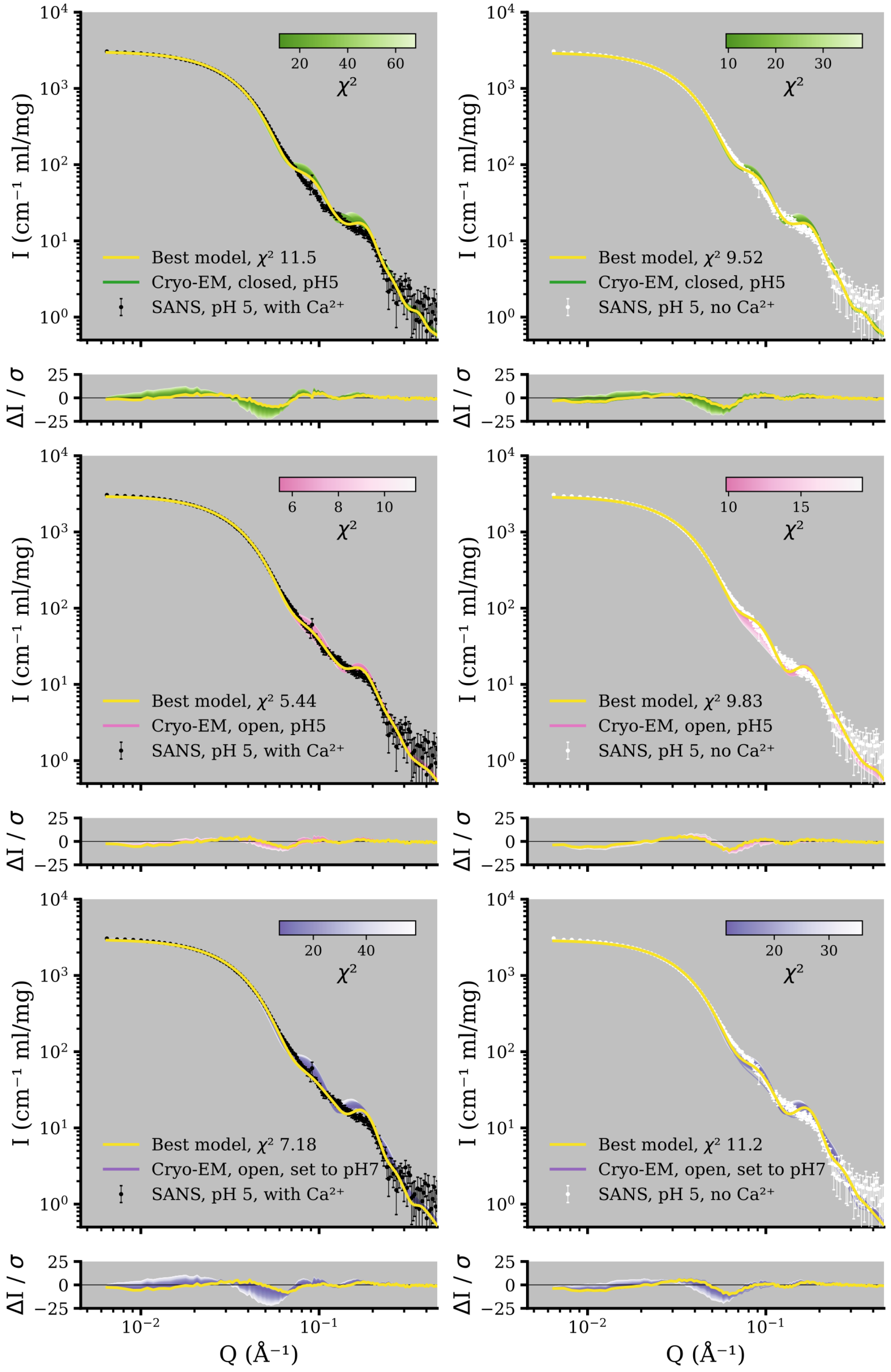


**Figure S14: Comparison of simulation snapshots to experimental SANS profiles.** Theoretical curves from simulation snapshots taken every 10 ns through the trajectories, compared to the experimental SANS profiles from DeCLIC at pH 5 in the presence (left column, black scattering data) and absence (right column, white scattering data) of calcium. Each row shows theoretical curves from one simulation condition; the first row shows the protonated simulations of the pH 5 closed cryo-EM structure in shades of green, the middle row shows the protonated simulations of the pH 5 open cryo-EM structure in shades of pink, the last row shows the deprotonated simulations of the pH 5 open cryo-EM structure in shades of lavender. The reduced $\chi^2$-value of each theoretical curve is indicated by its shade, with stronger color indicating a better goodness of fit, and the error weighted residual to the experimental curve shown in the lower panels of each row. The snapshot from each simulation condition with the best goodness of fit to the experimental data to which it is compared is shown in yellow.

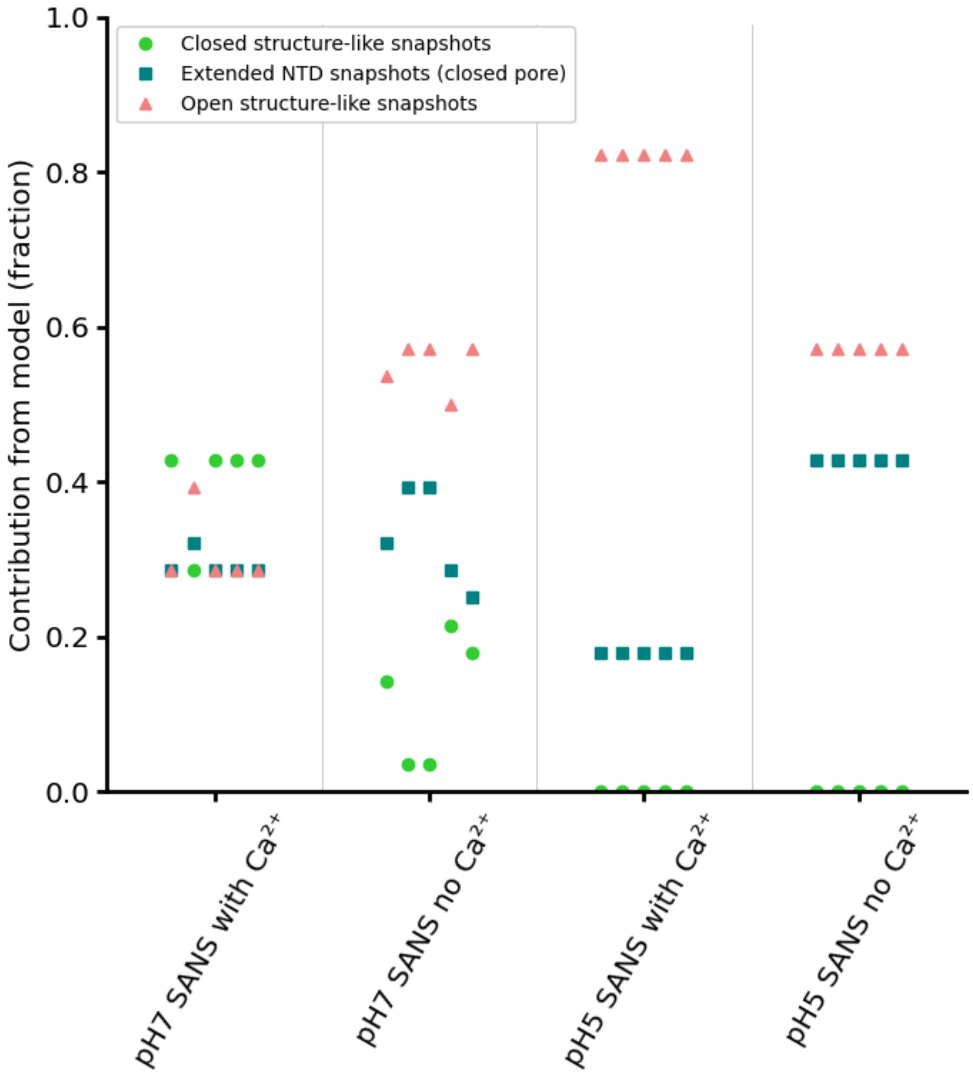


**Figure S15: Contributions to optimized ensembles fits to SANS data from DeCLIC** from five runs of the Gajoe genetic algorithm for each SANS dataset. The snapshots were visually inspected and as categorized as closed structure-like snapshots, open structure-like snapshots, and extended NTD snapshots, followed by contribution calculated as the number of snapshots from each respective category divided by the total number of snapshots in the ensemble. Markers vertically in line with each other are from the same ensemble. All repeats to the same SANS data had similar goodness of fit, with some minor variation depending on the composition of the ensembles. Goodness of fit ranges: $3.096 \leq \chi^2 \leq 3.173$ for pH7 with calcium, $3.329 \leq \chi^2 \leq 3.356$ for pH7 without calcium, $3.250 \leq \chi^2 \leq 3.286$ for pH5 with calcium, and $\chi^2 = 3.928$ for pH5 without calcium.

**Table S1:** Cryo-EM data collection and refinement statistics

| *Data Collection* | **With $Ca^{2+}$** | **No $Ca^{2+}$** |
|---|---|---|
| Microscope | FEI Titan Krios | |
| Magnification | 105,000 | |
| Voltage (kV) | 300 | |
| Electron exposure ($e^-$/$Å^2$) | $\sim 43$ | |
| Defocus Range (µm) | -1.4 – -3.0 | -1.6 – -3.0 |
| Pixel Size (Å) | 0.8617 | |
| Symmetry Imposed | C5 | |
| Number of Images | 26,259 | 26,757 |
| Particles Picked | $\sim 4$ mil | $\sim 3,5$ mil |
| Particles Refined | | |
| Open (EMDB) | 27,419 (50745) | 11,612 (50746) |
| Closed/Disordered (EMDB) | 22,391 (50744) | 169,422 (50747) |
| *Refinement* | | |
| Resolution (Å) | | |
| Open (PDB) | 2.9 (9FTH) | 3.5 (9FTI) |
| Closed/Disordered (PDB) | 3.1 (9FTG) | 4.0 (9FTJ) |
| FSC Threshold | 0.143 | |
| Map sharpening B-factor | | |
| Open | -65 | -120 |
| Closed/Disordered | -114 | -196 |

**Table S2:** Cryo-EM model refinement statistics

| | With $Ca^{2+}$ | | No $Ca^{2+}$ | |
|---|---|---|---|---|
| | **Open** (9FTH) | **Closed** (9FTG) | **Open** (9FTI) | **Disordered** (9FTJ) |
| Model Composition | | | | |
| Non-hydrogen Protein atoms | 22,780 | 21,205 | 19,945 | 12,060 |
| Protein Residues | 3010 | 2795 | 2780 | 1570 |
| Ligands | 0 | 5 | 0 | 0 |
| B-factor ($Å^2$) | 120.1 | 61.8 | 76.12 | 56.87 |
| RMSD | | | | |
| Bond Lengths (Å) | 0.002 | 0.003 | 0.005 | 0.007 |
| Bond Angles (°) | 0.461 | 0.487 | 0.597 | 0.644 |
| Validation | | | | |
| Molprobity Score | 1.21 | 1.61 | 1.69 | 1.69 |
| Clashscore | 3.48 | 4.22 | 5.62 | 6.09 |
| Poor Rotamers (%) | 0 | 0 | 0 | 0 |
| Ramachandran Plot | | | | |
| Favored (%) | 97.67 | 93.94 | 94.30 | 94.87 |
| Allowed (%) | 2.33 | 6.06 | 5.70 | 5.13 |
| Outliers (%) | 0 | 0 | 0 | 0 |

**Table S3:** Unmodeled residues in cryo-EM structures

| | With $Ca^{2+}$ | | No $Ca^{2+}$ | |
|---|---|---|---|---|
| | **Open** (9FTH) | **Closed** (9FTG) | **Open** (9FTI) | **Disordered** (9FTJ) |
| NTD1 | 28 – 34 | 28 – 37 | 28 – 38 | 28 – 200 |
| | | 58 – 66 | 54 – 66 | |
| | | 131 – 133 | | |
| | | | 138 – 145 | |
| | | 173 – 188 | 181 – 187 | |
| NTD2 | | 263 – 267 | | 201 – 325 |
| | | 291 – 293 | 290 – 298 | |
| | | 313 – 318 | 314 – 318 | |
| C-terminus | 637 – 642 | 639 – 642 | 637 – 642 | 640 – 642 |
| Total | 12 | 56 | 59 | 301 |

**Table S4:** Summary of electrophysiology recordings in planar lipid bilayers

| | With $Ca^{2+}$ | | No $Ca^{2+}$ | |
|---|---|---|---|---|
| | **pH 7** | **pH 5** | **pH 7** | **pH 5** |
| Membranes recorded | 20 | 21 | 20 | 19 |
| Recording time (s) | 516 ± 24 | 435 ± 28 | 479 ± 28 | 444 ± 19 |
| Fraction active (%) | 0.12 ± 0.08 | 2.1 ± 2.1 | 1.3 ± 0.9 | 8 ± 3.1 |
| Events per recording | 2.25 ± 1.53 | 1.48 ± 1.48 | 3.00 ± 1.79 | 11.00 ± 3.75 |
| Total events | 45 | 31 | 60 | 209 |
| Event duration (s) | 1.47 ± 0.22 | 6.81 ± 1.64 | 2.78 ± 0.59 | 3.38 ± 0.50 |
| Total time (s) | 10,321 | 9,127 | 8,418 | 8,433 |
| Total activity (s) | 66 | 211 | 167 | 706 |
| Total fraction active (%) | 0.64 | 2.30 | 2.00 | 8.00 |

**Table S5:** Sample details for SANS data collection

| | |
|---|---|
| Organism of origin | *Desulfofustis* sp. PB-SRB1 |
| Expression system | *Echerichia coli* |
| UniProt ID | V4JF97 |
| Extinction coefficient [$A_{280}$ 0.1%(w/v)] | 1.107 |
| Volume from structure (Å$^3$) | 495062 |
| Particle contrast from sequence and solvent constituents, $\Delta\bar{\rho}$ ($\lvert\rho_{protein} - \rho_{solvent}\rvert$; $10^{10}$ cm$^{-2}$) | 3.94 ($\lvert 2.44 - 6.38 \rvert$) |
| M from chemical composition (kDa) | 341.7 |

**Table S6:** Summary of SANS data collection parameters

| | |
|---|---|
| Instrument | ILL D22 |
| Wavelength (Å) | 6 |
| Pathlength (cm) | 0.1 |
| Detector distances (m) | 1.4m and 8m |
| $Q$ measurement range (Å$^{-1}$) | 0.006 – 0.730 |
| Absolute scaling method | Incident beam flux |
| Normalization | Divided by concentration |
| Exposure time (aggregate time) | |
| With $Ca^{2+}$ | 161 x 30s (80.5 min) |
| No $Ca^{2+}$ | 210 x 30s (105 min) |
| Column | Superdex 200 Increase 10/300 |
| Sample temperature ($^o$C) | 10 |
| Loading concentration | 5.5 mg/ml |
| Injection volume | 250 µl |
| Flow rate (ml/min) | 0.2, 0.01, 0 |
| Average concentration (mg/ml) in combined data frames | |
| With $Ca^{2+}$ | 1.4 |
| No $Ca^{2+}$ | 1.35 |
| Solvent | |
| With $Ca^{2+}$ | $D_2O$, 150 mM NaCl, 20 mM Citrate·HCl, 10 mM $CaCl_2$, 0.5 mM d-DDM |
| No $Ca^{2+}$ | $D_2O$, 150 mM NaCl, 20 mM Citrate·HCl, 10 mM EDTA, 0.5 mM d-DDM |

**Table S7:** Summary of structural parameters calculated from SANS data

| | With $Ca^{2+}$ | No $Ca^{2+}$ |
|---|---|---|
| Guinier analysis | | |
| $R_g$ (Å) | 50.7 ± 0.2 | 51.6 ± 0.2 |
| $Q_{min}$ (Å$^{-1}$) | 0.010 | 0.010 |
| $Q_{max}$ (Å$^{-1}$), ($QR_g$ max) | 0.035 (1.7) | 0.035 (1.7) |
| Coefficient of correlation $R^2$ | 0.9998 | 0.9999 |
| $P(r)$ analysis | | |
| $R_g$ (Å) | 50.21 ± 0.02 | 51.39 ± 0.06 |
| $d_{max}$ (Å) | 142.48 ± 0.78 | 149.95 ± 2.57 |
| $q$ range (Å$^{-1}$) | 0.0074 - 0.7155 | 0.0074 - 0.7155 |
| $\chi^2$ | 0.59 | 0.45 |
| PepsiSANS | | |
| $R_g$ (Å) | 49.7 | 51.2 |

**Table S8:** Summary of modeling using protein structures and structure-based models

| Structures | pH5 closed | pH5 open | |
|---|---|---|---|
| PepsiSANS | | | |
| Predicted $R_g$ (Å) | 51.1 | 49.1 | |
| SANS with $Ca^{2+}$ $\chi^2$ | 47.20 | 6.82 | |
| SANS no $Ca^{2+}$ $\chi^2$ | 28.15 | 16.30 | |
| MD-simulation models | pH 5 closed | pH 5 open | pH 5 open |
| Protonation state | Protonated | Protonated | Deprotonated |
| Equilibration time (ns) | 12 | 15 | 12 |
| Simulation time (ns/replica) | 1000 | 1000 | 1000 |
| Replicas (nr) | 4 | 4 | 4 |
| $\chi^2$ range (PepsiSANS) | | | |
| SANS with $Ca^{2+}$ | 11.50 - 68.34 | 5.44 - 11.35 | 7.18 - 58.63 |
| SANS no $Ca^{2+}$ | 9.52 - 38.29 | 9.83 - 19.24 | 11.20 - 36.10 |

**Table S9:** Summary of ensemble optimization results.

| **Ensemble optimization for SANS pH 5 with $Ca^{2+}$** | | | | | |
|---|---|---|---|---|---|
| $\chi^2$ | 3.250 | 3.250 | 3.250 | 3.250 | 3.286 |
| Nr of closed snapshots | 0 | 0 | 0 | 0 | 0 |
| Nr of extended NTD snapshots | 5 | 5 | 5 | 5 | 5 |
| Nr of open snapshots | 23 | 23 | 23 | 23 | 23 |
| **Ensemble optimization for SANS pH 5 without $Ca^{2+}$** | | | | | |
| $\chi^2$ | 3.928 | 3.928 | 3.928 | 3.928 | 3.928 |
| Nr of closed snapshots | 0 | 0 | 0 | 0 | 0 |
| Nr of extended NTD snapshots | 12 | 12 | 12 | 12 | 12 |
| Nr of open snapshots | 16 | 16 | 16 | 16 | 16 |
| **Ensemble optimization for SANS pH 7 with $Ca^{2+}$** | | | | | |
| $\chi^2$ | 3.125 | 3.173 | 3.126 | 3.096 | 3.096 |
| Nr of closed snapshots | 12 | 8 | 12 | 12 | 12 |
| Nr of extended NTD snapshots | 8 | 9 | 8 | 8 | 8 |
| Nr of open snapshots | 8 | 11 | 8 | 8 | 8 |
| **Ensemble optimization for SANS pH 7 without $Ca^{2+}$** | | | | | |
| $\chi^2$ | 3.352 | 3.354 | 3.354 | 3.329 | 3.356 |
| Nr of closed snapshots | 4 | 1 | 1 | 6 | 5 |
| Nr of extended NTD snapshots | 9 | 11 | 11 | 8 | 7 |
| Nr of open snapshots | 15 | 16 | 16 | 14 | 16 |

**Table S10:** Summary of software and equations employed for SANS data reduction, analysis, and interpretation.

| | |
|---|---|
| Momentum transfer | Q=(4π/λ)sin($\theta$) |
| SANS data reduction | Grasp v. 10.02 [71, 72] |
| Extinction coefficient estimate | ProtParam [73] |
| Guinier equation | $\ln(I(Q)) = \ln(I(0)) - \frac{R_g^2}{3}Q^2$ |
| Calculation of $\rho$ | $\rho = (\sum_{i=1}^{N} b_i)/V$ |
| Calculation of $\bar{\nu}$ | $\bar{\nu} = V/M_{aa}$ |
| Protein volume estimation | $^3$V: Voss Volume Voxelator [78] |
| Relationship between Q and distance | $d = 2\pi/$Q |
| $P(r)$ analysis | BayesApp [74] *via* web server (https://somo.chem.utk.edu/bayesapp/) |
| $P(r)$ from structure | CaPP [75] (https://github.com/Niels-Bohr-Institute -XNS-StructBiophys/CaPP) |
| Atomic structure modelling | PepsiSANS v. 3.0 [76] |
| Simulation system set-up | Charmm-GUI membrame builder [61, 62] |
| Molecular dynamics simulations | Gromacs v. 2021.3 [65] |
| Theoretical $R_g$ | PepsiSANS v. 3.0 [76] |
| Ensemble optimization | Eom (Gajoe) v. 2.1 [35, 36] |
| Hydrogen-deuterium exchange | Psx [79] |
| $pK_a$ estimation | Propka3 [31, 32] |
| Three-dimensional graphic model representation | Vmd [66], UCSF ChimeraX [60] |
| Plots | Matplotlib [80] |
| Pore profiles | Chap [67] |
